# The AGEF-1/ARF-1 GTPase/AP-1 trafficking pathway differentially regulates LIN-12/Notch signaling in a tissue specific manner in *C. elegans*

**DOI:** 10.64898/2026.04.17.719071

**Authors:** Tatsuya Kato, Clare FitzPatrick, Soroosh Siyoofi, Haojun Zhu, Edouarda Taguedong, Olga Skorobogata, Christian E. Rocheleau

## Abstract

LIN-12/Notch signaling regulates *C. elegans* vulval development via cell fate specifications in the gonad and epidermis. In the somatic gonad LIN-12/Notch activity specifies the anchor cell (AC) versus ventral uterine cell (VU) fates, with VU receiving more signal. The AC secretes epidermal growth factor (EGF) which induces the underlying vulval precursor cells (VPCs) to adopt vulval fates. In the VPCs the secondary vulval fates are specified by LIN-12/Notch activity. We previously reported that the AGEF-1, an Arf GEF homologous to ArfGEF1 and ArfGEF2, the ARF-1 GTPase, and the adaptor protein complex 1 (AP-1) inhibit LET-23/EGF receptor (EGFR) signaling in the VPCs by antagonizing LET-23/EGFR basolateral localization. Here we report that AGEF-1, ARF-1 and AP-1 regulate LIN-12/Notch signaling during somatic gonad and vulval development. The *lin-12(n302)* partial gain-of-function causes a potent Vulvaless phenotype due to a lack of AC specification. We demonstrate that loss of AGEF-1, ARF-1 or AP-1 restored the AC fate in *lin-12(n302)* animals, indicating that AGEF-1/ARF-1/AP-1 promotes LIN-12/Notch signaling in the somatic gonad. Interestingly, loss of AGEF-1, ARF-1 or AP-1 also induced ectopic vulval secondary fates in *lin-12(n302)* animals, indicating that AGEF-1/ARF-1/AP-1 inhibits LIN-12/Notch in the VPCs. Using a LIN-12/Notch biosensor we demonstrate that loss of UNC-101/AP-1μ results in decreased signaling in the VU cell and increased signaling in the VPCs that correspond with decreased expression levels of LIN-12/Notch and LAG-1/DSL ligand in the presumptive AC and VU while also causing increased apical localization of LIN-12/Notch in the VPCs. We hypothesize that the differential regulation of LIN-12/Notch signaling could reflect different trafficking pathways in epithelial cells (VPCs) versus non-epithelial cells (AC and VU). Our results indicate that the AGEF-1/ARF-1/AP-1 trafficking pathway maintains the VPC cell fate patterning by limiting both LET-23/EGFR and LIN-12/Notch signaling.

**Author summary:** Cell signaling and membrane trafficking are highly interconnected processes whereby membrane trafficking can regulate signal transduction pathways and vice versa. We previously demonstrated that the ARF-1 GTPase, the downstream AP-1 clathrin adaptor and upstream activator AGEF-1 antagonize the membrane trafficking of the Epidermal Growth Factor Receptor (EGFR) and hence signaling during *C. elegans* vulva induction. Strong loss of the ARF-1 GTPase pathway resulted in ectopic vulval induction. Here we demonstrate that the ARF-1 GTPase pathway differentially regulates Notch signaling to regulate vulva induction. In the somatic gonad it promotes Notch signaling to regulate the specification of the anchor cell which secretes the inductive signal. In the vulva precursor cells, the ARF-1 GTPase pathway antagonizes Notch signaling which cooperates with EGFR signaling to induce the vulval cell fates. We hypothesize that the differential regulation of Notch signaling by the ARF-1 GTPase pathway could be a result of more complex membrane trafficking pathways in polarized epithelial cells (vulva precursors) versus non-epithelial cells in the developing somatic gonad. Thus, the AGEF-1/ARF-1/AP-1 antagonizes both EGFR and Notch signaling in ensuring that only three of the six vulval precursor cells adopt are induced.

## Introduction

Membrane trafficking and signal transduction are tightly interconnected processes. The secretory pathway ensures proper delivery of transmembrane receptors and ligands to the plasma membrane and facilitates the release of soluble ligands into the extracellular space [1]. Endocytosis, in turn, enables receptor internalization, directing them either toward lysosomal degradation or recycling back to the membrane [2]. Together, these trafficking routes dynamically regulate the availability of ligands and receptors, thereby modulating signaling capacity.

Polarized cells such as neurons and epithelial cells require precise targeting of receptors and ligands to specific domains like axons versus dendrites or apical versus basolateral membranes [3]. Sorting decisions at the trans-Golgi network (TGN) and recycling endosomes are thus crucial for proper cellular function and overall organismal health. Imbalances in receptor sorting can lead to decreased or increased signaling and have pathological consequences.

The AP-1 clathrin adaptor complex is an important component of clathrin-mediated trafficking of cargo between the TGN, endosomes and the plasma membrane [3, 4]. AP-1 is a heterotetramer composed of γ1, α1, μ1, and β1 subunits. AP-1 is recruited to membranes in part by class I and II Arf GTPases. AP-1 recruits clathrin to aid in vesicle formation and AP-1 recruits cargo proteins via tyrosine and dileucine motifs into newly formed vesicles. Disruption of AP-1 leads to sorting of proteins to the inappropriate membrane domains and thus impacts cellular polarity and signal transduction. Loss of function mutations in *AP1S1* that codes for the σ1A subunit and *AP1B1* that codes for the β1 subunit are found to cause IDEDNIK (Intellectual Disability, Enteropathy, Deafness, Neuropathy, Ichthyosis, and Keratoderma) a rare autosomal recessive genetic disorder formerly known as MEDNIK [5, 6].

The AP-1 complex is conserved in the nematode *Caenorhabditis elegans* comprised of APG-1 γ1, APS-1σ1, APB-1 β1 (this subunit is shared with the AP-2 complex) and two proteins UNC-101 and APM-1 comprise the μ1 subunits [7, 8]. All but UNC-101 are essential for viability. Mutations in *unc-101* were identified as having locomotion defects, defects in dye uptake by sensory neurons and as suppressors of vulva induction phenotypes caused by mutations in *let-23/egfr* [9]. Loss of *unc-101* results in defective sorting of ciliary proteins and of dendritic receptors in neurons [10–12]. AP-1 is required for the maintenance of apicobasal polarity of the *C. elegans* intestinal cells [13–15]. RNAi knockdown of AP-1 components results in the mislocalization of the PAR-6 polarity protein as well as the apical and basal mislocalization of membrane proteins and expansion of the apical domains into the lateral membranes of the intestinal cells. We previously demonstrated that AP-1, the ARF-1 and ARF-5 GTPases as well as the AGEF-1 Arf GEF regulate the apical versus basal localization of LET-23/EGFR in the vulval precursor cells [16]. Disruption of AGEF-1/Arf GTPase/AP-1 mediated trafficking increased basolateral LET-23/EGFR localization as well as inducing ectopic vulva induction [7, 16].

Sequential signaling by the LET-23/EGFR and LIN-12/Notch are required for the induction and patterning of the *C. elegans* vulva [17, 18]. LIN-12/Notch functions in the developing somatic gonad to specify the anchor cell (AC) and ventral uterine precursor (VU) cell [19, 20]. Two equipotent α cells (Z1.ppp and Z4.aaa) express LIN-12/Notch and LAG-2/Delta ligand [21]. The cell that receives the highest amount of signal adopts the VU fate and the other cell the AC fate. Loss of LIN-12/Notch signaling results in both cells adopting the AC fate and increased signaling results in both cells adopting the VU fate [19]. The AC secretes LIN-3/Epidermal Growth Factor (EGF) which induces three of six multipotent epithelial cells, the vulval precursor cells (VPCs; P3.p-P8.p) to adopt vulval fates [22]. P6.p, the VPC most proximal to the LIN-3/EGF secreting AC is induced to adopt a primary fate by activation of LET-23/EGFR signaling. Graded LIN-3/EGF signaling as well as lateral LIN-12/Notch signaling induces the adjacent P5.p and P7.p cells to adopt the secondary fate [23]. LIN-12/Notch signaling is both necessary and sufficient to induce the secondary vulval fates [19]. The uninduced VPCs (P3.p, P4.p, P8.p) take on the tertiary non-vulval fate. Aberrations to this network lead to observable developmental defects such as ectopic vulval inductions leading to a Multivulva (Muv) phenotype or reduced inductions causing a Vulvaless (Vul) phenotype [24].

In Drosophila, AP-1 has been shown to promote and antagonize Notch signaling in several tissues. For example, AP-1 antagonizes Notch signaling during sensory organ development. Loss of AP-1 causes partial Notch gain-of-function phenotypes that are dependent on the Notch regulator Sanpodo. AP-1 interacts with Sanpodo and regulates the localization of Notch and Sanpodo [25]. During eye development, AP-1 promotes Notch signaling as AP-1 knockdown results in two R8 cells in the ommatidia suggesting a defect in Notch mediated lateral inhibition [26]. Reduced Notch levels in AP-1 mutants are attributed to defects in secretion of Scabrous which functions to stabilize Notch at the plasma membrane to prevent its endocytosis and lysosomal degradation. Thus, AP-1 differentially regulates Notch signaling in different tissues that may be dependent on different Notch trafficking regulators.

While AP-1 and the late endosomal RAB-7 GTPase regulate LET-23/EGFR trafficking and antagonize LET-23/EGFR signaling during vulva development, we found that simultaneous loss of UNC-101 and RAB-7 resulted in ectopic vulval inductions with characteristics of secondary vulval fates rather than primary suggesting a role in regulation of Notch signaling [27]. Here we demonstrate that AP-1, ARF-1 and AGEF-1 promote LIN-12/Notch signaling during the VU/AC decision in the somatic gonad but antagonize LIN-12/Notch signaling in the VPCs. Loss of *unc-101, arf-1* or *agef-1* potently suppresses the Vul phenotype of a weak activating allele of LIN-12/Notch restoring the specification of the AC. At the same time loss of *unc-101*, *arf-1* or *agef-1* enhances the ectopic induction of secondary fates in the VPCs. We demonstrate that in the absence *of unc-101*, there is a decrease in LIN-12/Notch signaling activity in the VU cell that correlated with decreased levels of LIN-12/Notch and its ligand LAG-2/DSL. While in the VPCs, LIN-12/Notch signaling is increased in the presumptive secondary vulval cells of *unc-101* mutants and there is an increase localization of LIN-12 on the apical membrane. *C. elegans* lack obvious homologs of Drosophila Sanpodo and Scabrous that could explain differences in signaling. Rather we hypothesize that this difference could reflect that the VPCs are epithelial cells and the AC is not. Thus, in the VPCs, the AGEF-1/ARF-1/AP-1 pathway antagonizes both LET-23/EGFR and LIN-12/Notch-mediated vulva induction to ensure that only three VPCs are induced.

## Results

### AP-1, ARF-1 and AGEF-1 are required for the Vul phenotype of a weak *lin-12/Notch* hypermorph

We previously found that an *unc-101(sy108); rab-7(ok511)* double mutant resulted in ectopic vulva induction in which the ectopically induced vulval cells appeared to adopt a secondary vulval fate suggesting that one or both trafficking proteins regulate LIN-12/Notch signaling [27]. To investigate the role of RAB-7 and AP-1 in regulation of LIN-12/Notch signaling during *C. elegans* vulval development, we analyzed effects of the *rab-7(ok511)* and *unc-101(sy108)* null alleles [9, 27] on the *lin-12(n302)* Notch receptor mutant background. In wild-type hermaphrodites, proper signaling results in reproducible induction of the P5.p, P6.p and P7.p VPCs with a stereotypical vulval morphology at the mid-fourth larval stage (mid L4; Fig. 1A). By contrast, *lin-12(n302)* is a LIN-12/Notch receptor weak hypermorph with a strong Vul phenotype due to the absence of the AC but a failure to induce ectopic secondary fates as seen in strong *lin-12* hypermorphs, making it amenable to enhancement or suppression [19, 28] (Fig. 1B). In this Notch-sensitized background, we found that *rab-7(ok511)* mildly increased vulva induction of *lin-12(n302)* from an average of 0.07 VPCs induced (*n*=22) to 0.42 (*n*=23, unpaired t test *P=0.0178*). In contrast, *unc-101(sy108)* strongly suppressed the Vul phenotype and rescued vulval development (Fig. 1C), indicated by a significant increase in the average number of induced VPCs (Fig. 1G). Notably, the percentage of animals with wild-type vulval morphology increased from 0% (0/42) in the *lin-12(n302)* background to 26% (17/65) in *unc-101(sy108); lin-12(n302)* (Fig. 1C,G). These results identify *unc-101(sy108)* as a strong suppressor of the *lin-12(n302)* Vul phenotype, pointing to a role for AP-1 in regulating LIN-12/Notch signaling during vulval development.

**Fig. 1.**
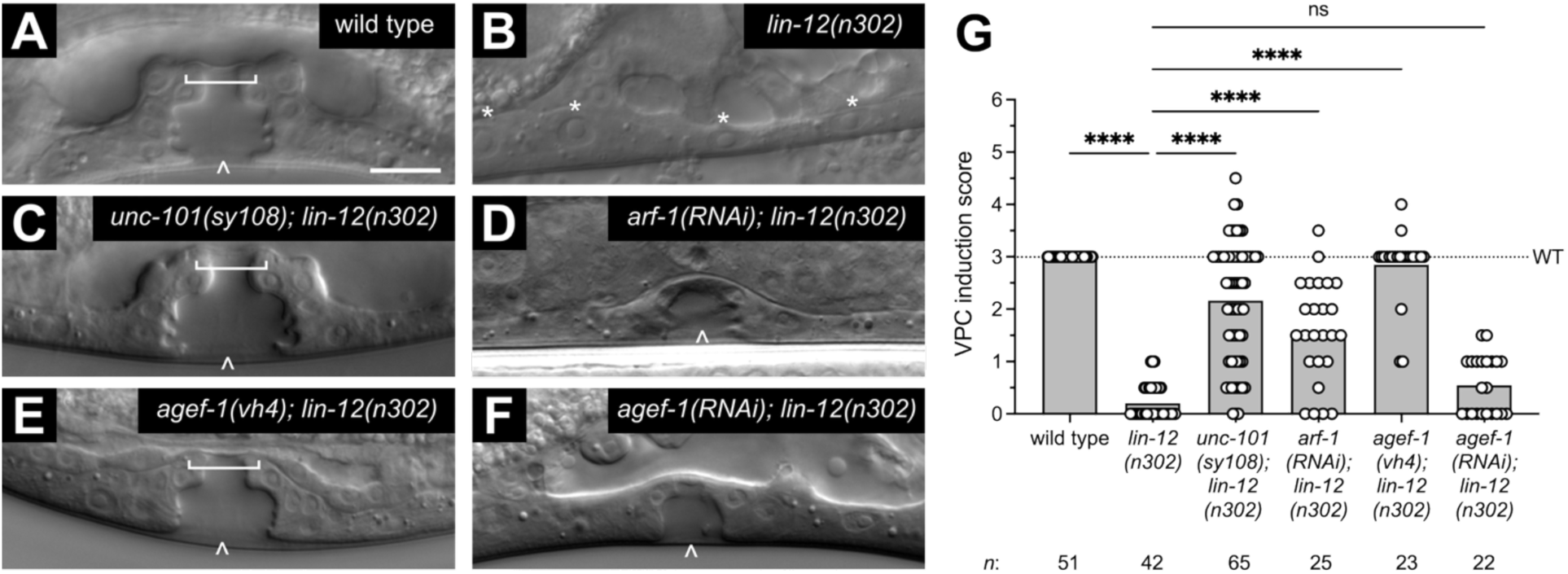
Loss of *unc-101, arf-1* and *agef-1* suppress the *lin-12(n302)* Vul phenotype. (A-F) Differential interference contrast (DIC) images of developing vulvae of wild type, *lin-12(n302)*, *unc-101(sy108); lin-12(n302)*, *arf-1(RNAi); lin-12(n302)*, *agef-1(vh4); lin-12(n302)* and *agef-1(RNAi); lin-12(n302)* at the mid-fourth larval stage (mid L4). Carets in (A,C,D,E,F) denote vulval lumens. Brackets in (A,C,E) denote the utse. Asterisks in (B) denote uninduced VPC daughter cell nuclei. (G) VPC induction scores of wild type, *lin-12(n302)*, *unc-101(sy108); lin-12(n302)*, *arf-1(RNAi); lin-12(n302)*, *agef-1(vh4); lin-12(n302)* and *agef-1(RNAi); lin-12(n302)* at mid L4. Bars represent average VPC induction score, and dots represent VPC induction scores of individual larvae. Dotted line represents the wild-type VPC induction score of three. See “VPC induction scoring” in Materials and methods for details. One-way ANOVA. ns, not significant, ****P<0.0001. Scale bar, 10µm.

Since AP-1 negatively regulates LET-23/EGFR signaling with ARF-1 and AGEF-1 during vulval development [16], we tested if ARF-1 and AGEF-1 also regulate LIN-12/Notch signaling. We found that *arf-1(RNAi)* suppressed the *lin-12(n302)* Vul (Figure 1D,G), as did the *agef-1(vh4)* missense allele [16] (Fig. 1E,G). By contrast, *agef-1(RNAi)* led to a slight but non-significant suppression (Fig. 1F,G). This difference could be due to inefficient knockdown of *agef-1* by RNAi or a reflection of recent data demonstrating that the *agef-1(vh4)* allele behaves as an activating allele in some contexts as opposed to as a hypomorph [29]. These results suggest that AP-1 functions with ARF-1 and AGEF-1 to regulate LIN-12/Notch signaling during vulval development.

We tested *unc-101(sy108)* in the context of two additional *lin-12* alleles to further assess *unc-101*’s role in LIN-12/Notch signaling. The *lin-12(n920)* strong hypermorph induces ectopic secondary fate inductions in spite of lacking an AC, thus exhibiting a Multivulva phenotype [19] (Fig. S1A,E). The other tested allele *lin-12(n941)* is a LIN-12/Notch null which causes VPC fate mispatterning, leading to the protruded vulva phenotype [19] (Fig. S1C,F). *unc-101(sy108)* did not alter VPC induction scores or suppress the phenotypes of either allele (Fig. S1B,D,E,G,H), suggesting that AP-1 is not an essential downstream component of LIN-12/Notch signaling but rather plays a non-essential role modulating LIN-12/Notch signaling.

**Fig. S1.**
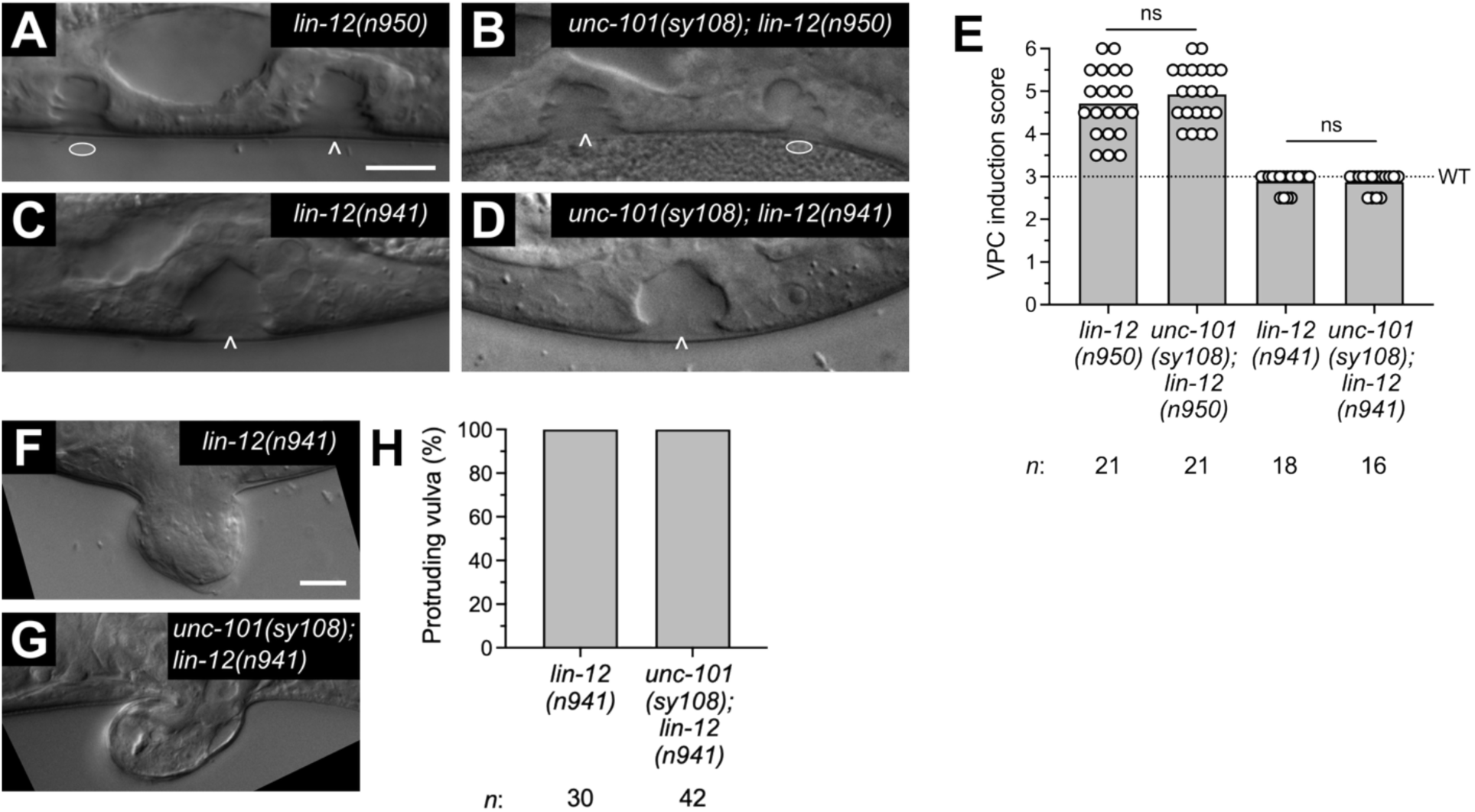
Loss of *unc-101* does not suppress or enhance strong activating or loss of function *lin-12* alleles. (A-D) DIC images of developing vulvae of *lin-12(n950)*, *unc-101(sy108); lin-12(n950), lin-12(n941),* and *unc-101(sy108); lin-12(n941)* at the mid-L4 stage. Carets in (A-D) denote vulval lumens resulting from induction of the P5.p, P6.p and P7.p VPCs. Ovals in (A,C) denote ectopic lumens. (E) VPC induction scores of *lin-12(n950)*, *unc-101(sy108); lin-12(n950), lin-12(n941),* and *unc-101(sy108); lin-12(n941)* at mid L4. One-way ANOVA. ns, not significant. (F,G) DIC images the protruding vulva phenotype of *lin-12(n941)* and *unc-101(sy108); lin-12(n941)* adults which were 100% penetrant (H). Scale bars, 10µm.

### Loss of *unc-101* rescues vulval development of *lin-12(n302)* by restoring the anchor cell fate

When the vulval lumen forms in wild type, the AC fuses with the syncytial uterine seam cell, utse [30, 31], observable as a thin bridge of tissue above the lumen at the mid-fourth larval stage (brackets on Fig. 1A,C,E). The wild-type vulval morphology and the presence of the cytoplasmic bridge in *unc-101(sy108); lin-12(n302)* indicates restoration of the AC fate resulting from AP-1 loss of function (Fig. 1C). To confirm, we used an endogenous LAG-2/DSL ligand nuclear reporter [32] which marks the AC fate (Fig. 2A). While somatic gonad nuclear signal was absent in *lin-12(n302)* (Fig. 2B,D), it was present in >50% of *unc-101(sy108); lin-12(n302)* double mutants (Fig. 2C,D). This taken together with the presence of the AC-derived cytoplasmic bridge in *agef-1(vh4); lin-12(n302)* animals suggests that the AGEF-1/ARF-1/AP-1 pathway promotes LIN-12/Notch signaling during the VU/AC decision.

**Fig. 2.**
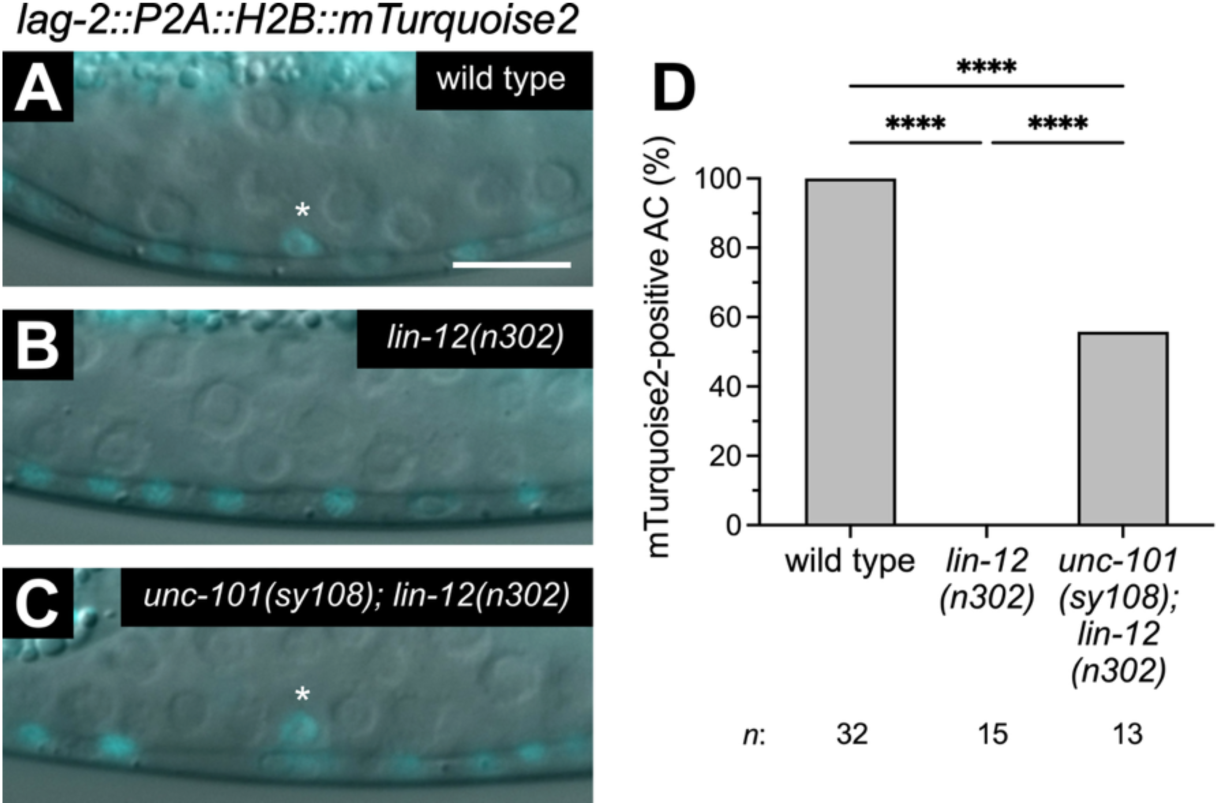
Loss of *unc-101* restores the anchor in *lin-12(n302)* mutants. (A-C) Merged DIC and epifluorescence images L2 stage larvae expressing the anchor cell (AC) reporter *lag-2::H2B::mTurquoise2, lag-2(bmd202),* in developing somatic gonads of wild type, *lin-12(n302)* and *unc-101(sy108); lin-12(n302)*. Asterisks in (A,C) denote anchor cell nuclei that is absent in *lin-12(n302)* (B). *lag-2::H2B::mTurquoise2* is also expressed in ventral cord neurons. (D) Percentage of mTurquoise2-positive AC nuclei in wild type, *lin-12(n302)* and *unc-101(sy108); lin*-*12(n302)*. One way ANOVA. ****P<0.0001. Scale bar, 10µm.

### AP-1 promotes LIN-12/Notch signaling in the somatic gonad

During specification of the AC from two bipotent somatic gonadal α cells, the α cell with high levels of LIN-12/Notch signaling commits to the VU fate while its neighbor receives less signal and becomes the AC [20, 33, 34]. The restored AC fate in *unc-101(sy108); lin-12(n302)* thus suggests reduced LIN-12/Notch signaling activity. To determine whether AP-1 promotes LIN-12/Notch signaling during the VU/AC decision, we analyzed the SALSA (Sensor Able to detect Lateral Signaling Activity) *in vivo* biosensor in wild-type and *unc-101(sy108)* larvae [35]. The SALSA biosensor consists of the C-terminus of LIN-12/Notch tagged with a TEV protease. Upon ligand binding and LIN-12 activation the C-terminus is released and enters the nucleus where it cleaves a GFP::RFP tagged histone resulting in the release of GFP into the cytoplasm. LIN-12/Notch signaling activity is measured by quantifying nuclear-to-cytoplasmic GFP localization change relative to histone-tagged nuclear RFP expression. Quantifying RFP:GFP nuclear fluorescence ratios with or without *unc-101(sy108)* indicated that loss of AP-1 correspondingly decreased LIN-12/Notch activity in the VU cell (Fig. 3A-C). No significant difference was seen in the AC; however, LIN-12/Notch signaling is lower in the AC than in the VU and would be less sensitive to perturbation of signaling.

**Fig. 3.**
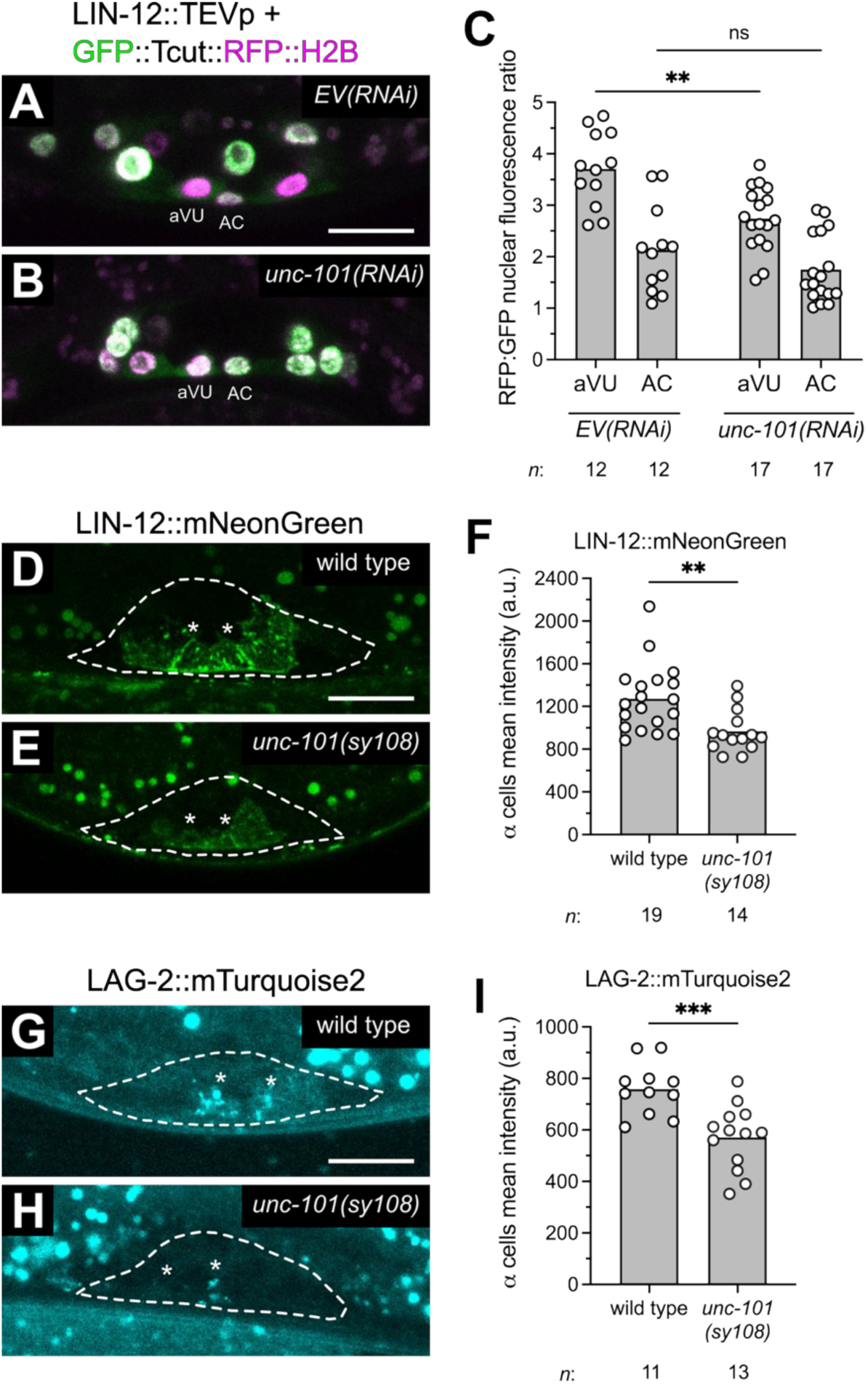
AP-1 promotes LIN-12/Notch signaling and LIN-12/Notch and LAG-2/DSL levels in the α cells of the somatic gonad. (A,B) Confocal images of the somatic gonads of *EV(RNAi)* and *unc-101(RNAi)* at L2 larvae expressing the LIN-12/Notch signaling biosensor *lin-12::TEVp + GFP::Tcut::RFP::H2B* (green and magenta). The presumptive a cell-derived ventral uterine cell (aVU) and anchor cell (AC) are labeled. (C) RFP:GFP nuclear fluorescence intensity ratios in *EV(RNAi)* and *unc-101(sy108)*. Multiple t-test. **P<0.01 (D,E) Confocal images of LIN-12::mNeonGreen, *lin-12(ljf33),* in the somatic gonads of wild type and *unc-101(sy108)* L2 larvae. (F) Mean fluorescence intensity of LIN-12::mNeonGreen in the a cells of wild type and *unc-101(sy108).* Unpaired t-test. ***P<0.001. (G,H) Confocal images of LAG-2::mTurquoise2, *lag-2(bmd204),* in the somatic gonads of wild type and *unc-101(sy108)* at L2 larvae. (I) Mean fluorescence intensity of LAG-2::mTurquoise2 in the a cells of wild type and *unc-101(sy108).* Unpaired t-test. **P<0.01. Dotted lines in (D,E,G,H) demarcate somatic gonad. Asterisks in (D,E,G,H) denote the a cells. Scale bars,10µm.

Since AP-1 regulates protein trafficking from the TGN to the plasma membrane and endosome recycling we analyzed the localization of endogenously-tagged LIN-12/Notch receptor and LAG-2/DSL ligand [32] in the α cells of wild-type and *unc-101(sy108)* larvae. LIN-12::mNeonGreen was expressed largely on the plasma membranes of the α cells (Fig. 3D) and LAG-2::mTurquoise2 displayed punctate expression (Fig. 3G). We found that *unc-101(sy108)* significantly decreased expression levels of both endogenous (Fig. 3B,C) and multicopy (Fig. S2A-C) reporters of LIN-12/Notch. Moreover, expression of the LAG-2/DSL ligand was also reduced (Fig. 3E,F). These results are consistent with AP-1 promoting trafficking of LIN-12/Notch and LAG-2/DSL to the plasma membranes of the α cells and thus may explain the reduced LIN-12/Notch signaling seen with SALSA in *unc-101(sy108)* mutants.

**Fig. S2.**
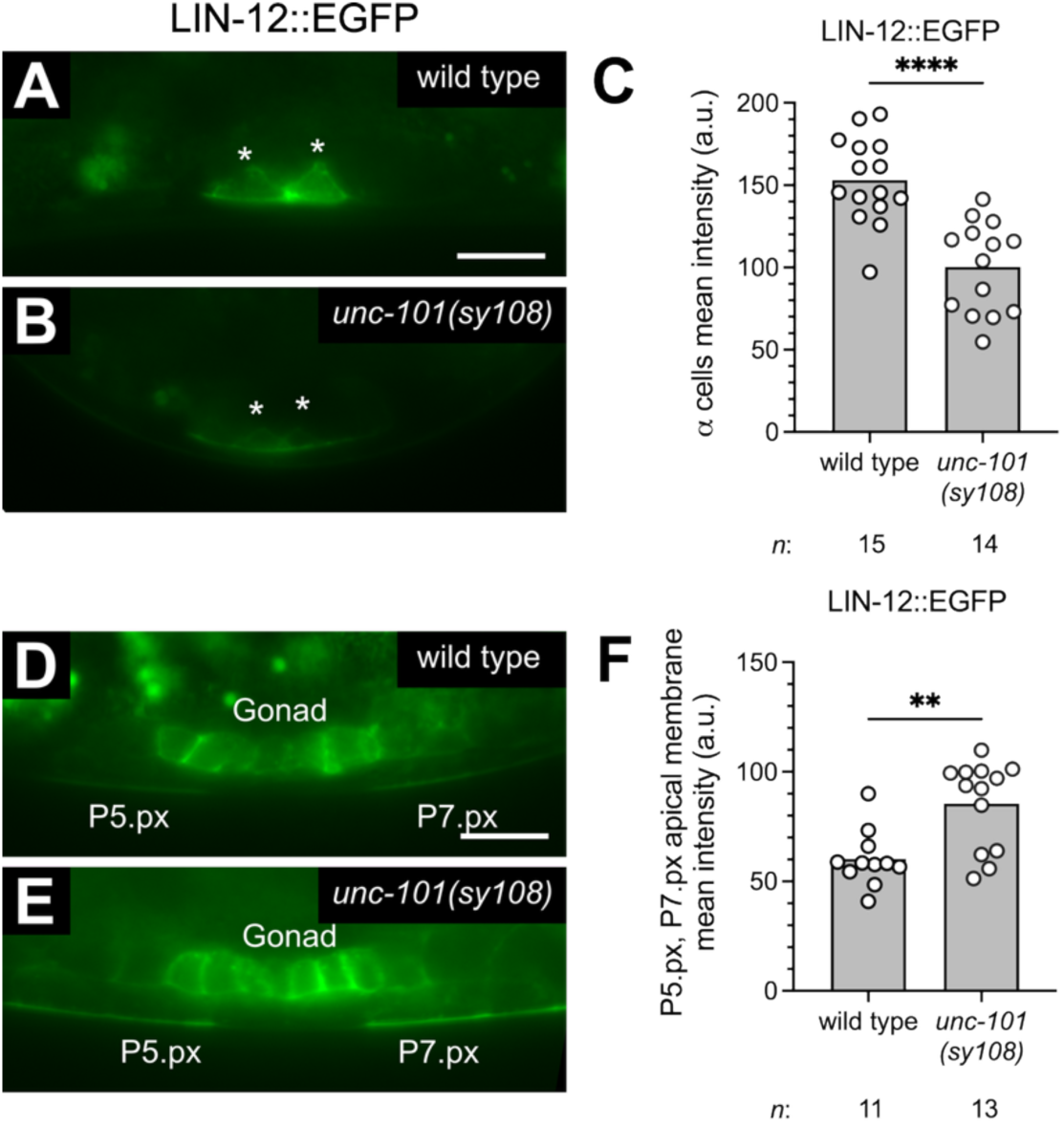
AP-1 regulates LIN-12/Notch localization in the somatic gonadal α cells and vulval precursor cells. (A,B) Confocal images of LIN-12::EGFP (*wgIs72*) in developing a cells of wild type and *unc-101(sy108)* at the second larval stage (L2). (C) Mean fluorescence intensity of the α cells in wild type and *unc-101(sy108).* Unpaired t-test. ****P<0.0001. (D,E) Confocal images of LIN-12::EGFP in VPCs of wild type and *unc-101(sy108)* in the VPC daughters (Pn.px). The somatic gonad, P5.px and P7.px are labeled. (F) Mean fluorescence intensities of P5.px, P7.px apical membranes in wild type and *unc-101(sy108)*. Unpaired t-test. **P<0.01.

### AP-1 inhibits LIN-12/Notch signaling in the vulval epithelium

While *unc-101(sy108), arf-1(RNAi),* and *agef-1(vh4)* each restored AC specification and hence, vulva induction in *lin-12(n302)*, we noted that some animals had greater than three VPCs induced suggesting an additional role in vulva induction (Fig. 1G). In fact, approximately 30% (19/65) of *unc-101(sy108); lin-12(n302)* animals displayed ectopic inductions of VPCs that are not normally induced (P3.p, P4.p or P8.p,) that were visible as ectopic lumens (Fig. 4A,B,E). Similar ectopic vulval inductions were observed in *arf-1(RNAi); lin-12(n302)* and *agef-1(vh4)*; *lin-12(n302)* animals (Fig. 4C-E). To determine whether these ectopic inductions are of primary or secondary fate and associated with LET-23/EGFR or LIN-12/Notch signaling upregulation, respectively, we analyzed transcriptional reporters of both pathways. In wild type, a primary fate reporter driven by the *egl-17/FGF* promoter [36] was expressed in P6.p with occasional expression in the presumptive secondaries P5.p and P7.p (Fig. S3A), while the secondary fate reporter driven by the *lst-5* promoter [37] was expressed exclusively in P5.p and P7.p (Fig. 4F). In *lin-12(n302)* larvae the expression of both reporters was mostly absent (Fig. 4G,I, Fig. S3B,D). In *unc-101(sy108); lin-12(n302)* double mutants the *lst-5* secondary fate reporter was expressed in all VPCs (Fig. 4H,I), while the *egl-17* primary fate reporter showed a wild-type pattern of expression (Fig. S3C,D), indicating that loss of AP-1 leads to induction of ectopic secondary fates in a LIN-12/Notch-sensitized background.

**Fig. 4.**
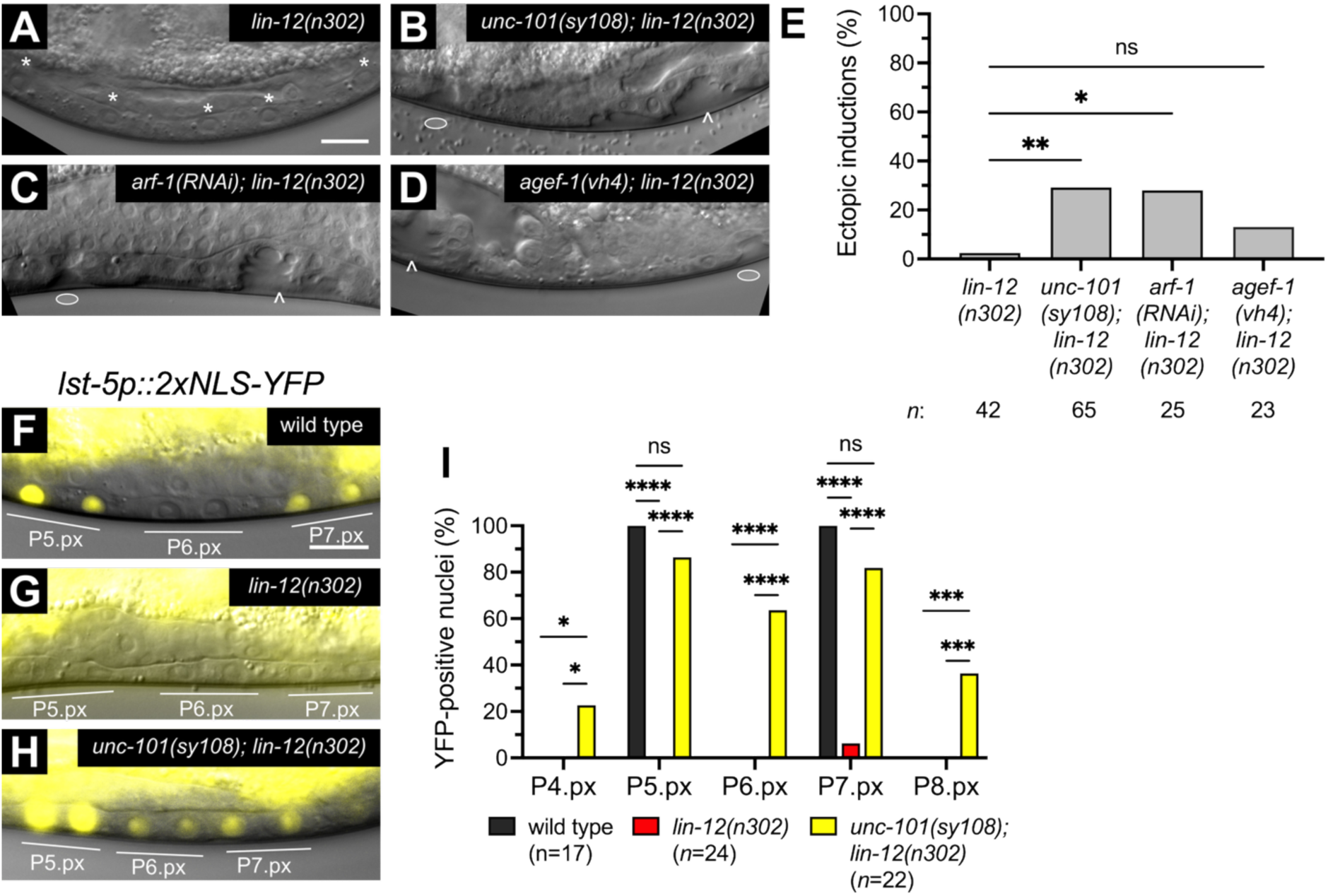
Loss of *unc-101* leads to induction of ectopic secondary fates in *lin-12(n302).* (A-D) DIC images of mid-L4 stage vulvae of *lin-12(n302)*, *unc-101(sy108); lin-12(n302)*, *arf-1(RNAi); lin-12(n302)* and *agef-1(vh4); lin-12(n302)*. Asterisks in (A) denote uninduced VPC daughter nuclei. Carets in (B-D) denote vulval lumens resulting from induction of the P5.p, P6.p and P7.p VPCs. Ovals in (B-D) denote vulval lumens resulting from ectopic inductions. (E) Percent ectopic inductions in *lin-12(n302)*, *unc-101(sy108); lin-12(n302)*, *arf-1(RNAi); lin-12(n302)* and *agef-1(vh4); lin-12(n302)* at mid L4. One-way ANOVA. ns, not significant, *P<0.05, **P<0.01. (F-H) Merged DIC and epifluorescence images of the secondary fate reporter *lst-5p::2xNLS-YFP* (yellow) in wild type, *lin-12(n302)* and *unc-101(sy108); lin-12(n302)* in the VPC daughters. (I) Percent YFP-positive nuclei in wild type, *lin-12(n302)* and *unc-101(sy108); lin-12(n302)*. Two-way ANOVA. ns, not significant, *P<0.05, ***P<0.001, ****P<0.0001. Scale bars, 10µm.

To determine if AP-1 regulates LIN-12/Notch signaling in a *lin-12(+)* background we tested the effects of the *unc-101(sy108)* mutation on the SALSA biosensor in the VPCs. We found a significant decrease in nuclear GFP (i.e., increased RFP:GFP ratio) in the secondary cells of *unc-101(sy108)* as compared to wild type (Fig. 5A-C). Thus, LIN-12/Notch signaling in the secondary cells was enhanced by loss of *unc-101* and suggesting that AP-1 antagonizes LIN-12/Notch signaling in the VPCs whereas it promotes signaling in the somatic gonad.

**Fig. 5.**
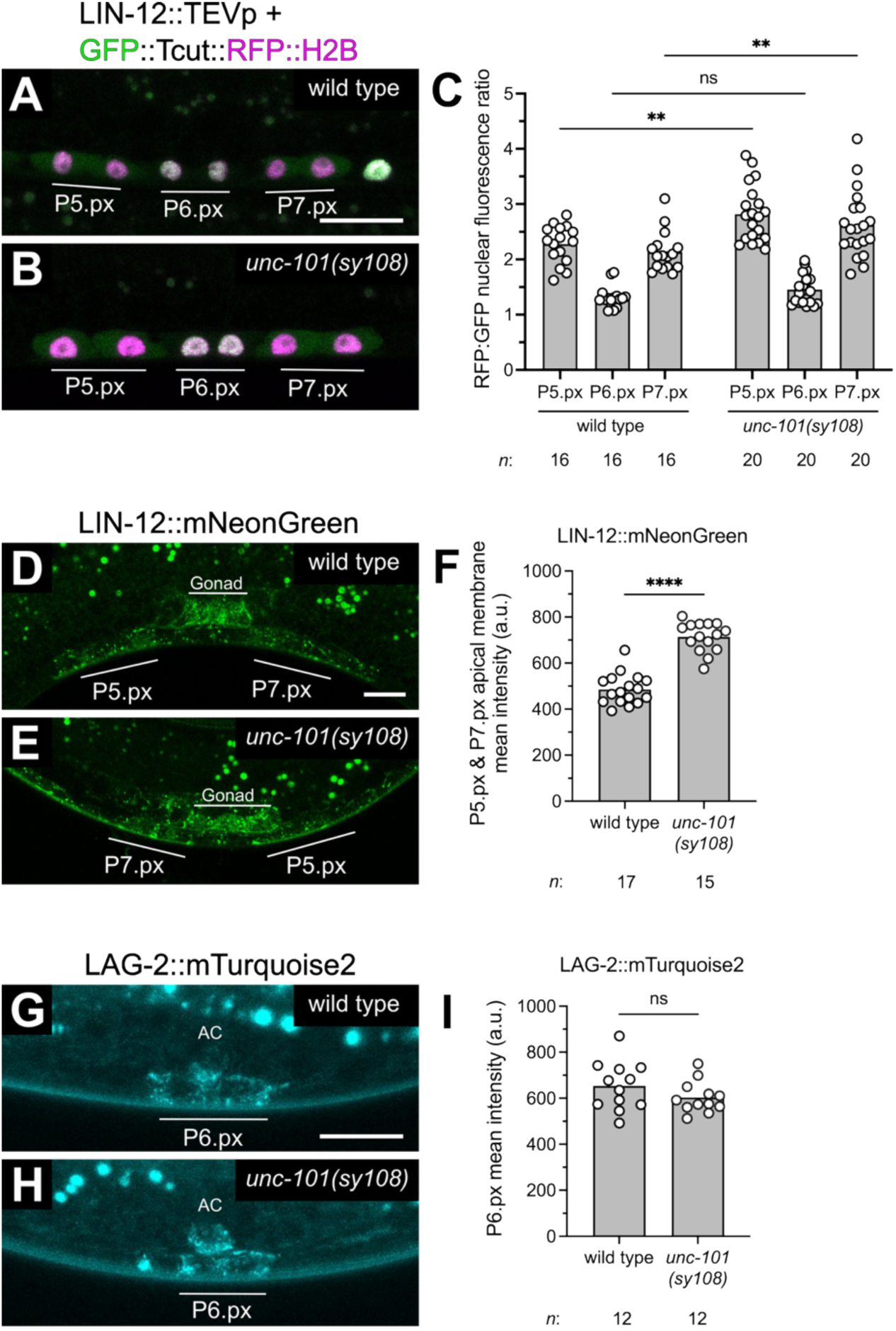
AP-1 inhibits LIN-12/Notch signaling and regulates LIN-12/Notch localization in the VPCs. (A,B) Confocal images of the VPC daughters expressing the LIN-12/Notch signaling biosensor *lin-12::TEVp + GFP::Tcut::RFP::H2B* (green and magenta) in wild-type and *unc-101(sy108)* larvae. (C) RFP:GFP nuclear fluorescence intensity ratios in wild type and *unc-101(sy108)*. Multiple t-test. ns, not significant, **P<0.01. (D,E) Confocal images of LIN-12::mNeonGreen, *lin-12(ljf33),* in the VPC daughters of wild type and *unc-101(sy108)*. (F) Mean fluorescence intensities of LIN-12::mNeonGreen on the apical membranes of P5.px and P7.px cells in wild type and *unc-101(sy108).* Unpaired t-test. ****P<0.0001. (G,H) Confocal images of LAG-2::mTurquoise2, *lag-2(bmd204),* in the P6.p daughter cells of wild type and *unc-101(sy108)*. Anchor cell (AC). (I) Mean fluorescence intensities of LAG-2::mTurquoise2 in the P6.px cells in wild type and *unc-101(sy108).* Unpaired t-test. ns, not significant. Scale bars,10µm.

To determine if AP-1 regulates the localization of the LIN-12/Notch receptor and LAG-2/DSL ligand in the VPCs we assessed their localization in *unc-101(sy108).* In the VPCs, LIN-12::mNeonGreen was expressed on the apical membranes of the secondary VPC daughter cells P5.px and P7.px (Fig. 5D), while LAG-2::mTurquoise2 displayed punctate expression near the P6.px membranes (Fig. 5G). In the *unc-101(sy108)* VPCs, we observed significantly increased apical membrane LIN-12/Notch expression (Fig. 5D-F; Fig. S2D-F). By contrast, there was no change in LAG-2/DSL ligand expression (Fig. 5G-I). Altogether these results indicate that AP-1 inhibits LIN-12/Notch signaling in the VPCs which contrasts with its function in the somatic gonad.

**Fig. S3.**
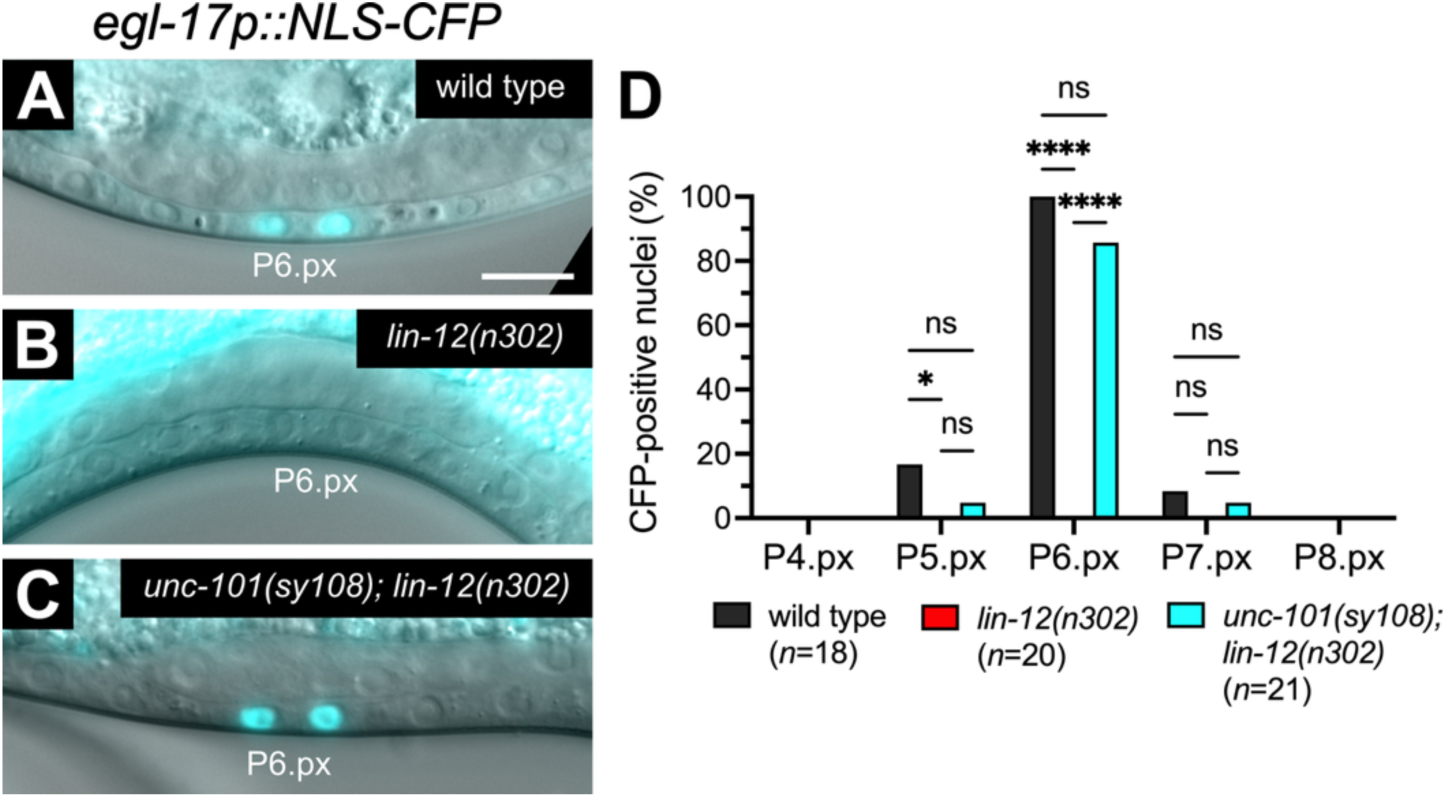
Loss of *unc-101* does not lead to ectopic primary fate inductions in *lin-12(n302).* (A-C) Merged DIC and epifluorescence images of the primary fate reporter *egl-17p::NLS-CFP* (cyan) in the VPC daughter cells of wild type, *lin-12(n302)* and *unc-101(sy108); lin-12(n302)*. (D) Percent CFP-positive nuclei in wild type, *lin-12(n302)* and *unc-101(sy108); lin-12(n302)*. Two-way ANOVA. ns, not significant, *P<0.05, ****P<0.0001. Scale bars, 10µm.

## Discussion

This study reveals differential regulation of LIN-12/Notch signaling by the AGEF-1/ARF-1/AP-1 trafficking pathway during vulval development (Fig. 6). Loss of the AP-1 μ1 subunit *unc-101/AP1M1* decreases LIN-12/Notch signaling activity in the somatic gonad while increasing it in the vulval epithelium which correlates with changes in LIN-12/Notch membrane expression. Although these expression changes do not appear to alter vulval development on its own, *unc-101(sy108)* restored the AC fate and enhanced ectopic secondary vulval fate inductions in the *lin-12(n302)* partial gain-of-function background, indicating AP-1’s roles in regulating LIN-12/Notch signaling during AC specification and secondary fate induction. AP-1 likely functions with the ARF-1 GTPase, its endosomal recruiter, and AGEF-1, an ARF-1 activator, to maintain steady-state levels of LIN-12/Notch signaling. Thus, the AGEF-1/ARF-1/AP-1 trafficking pathway plays an important role in VPC patterning by limiting both LET-23/EGFR and LIN-12/Notch signaling.

**Fig. 6.**
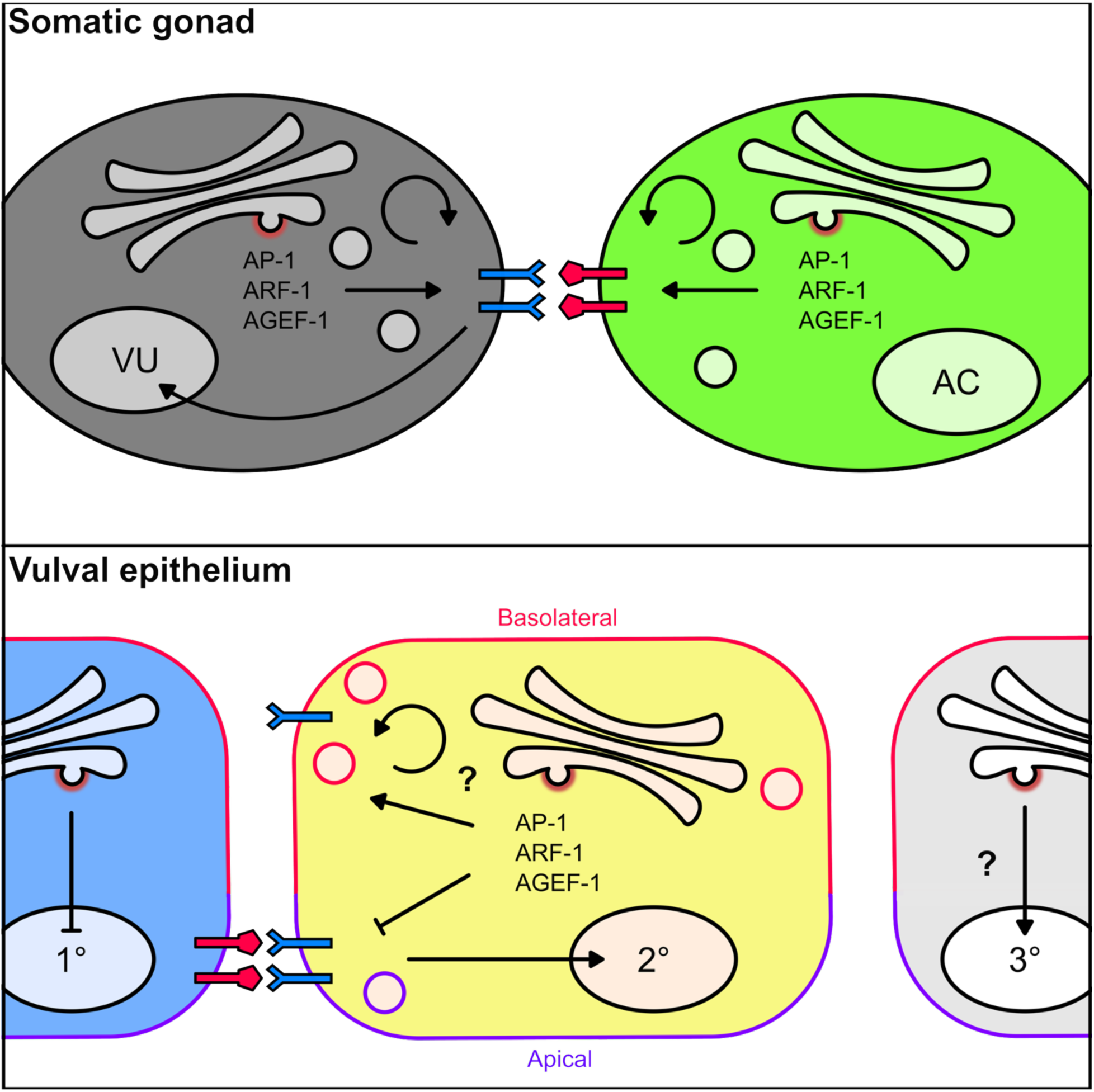
A model for differential LIN-12/Notch signaling regulation by AP-1 during *C. elegans* vulval development. (Top) In somatic gonadal α cells, AP-1 regulates sorting of LIN-12/Notch receptor (blue) and LAG-2/DSL ligand (red) between the trans-Golgi network and plasma membrane, affecting their secretion and/or recycling. AP-1 likely works together with ARF-1 and AGEF-1 to regulate receptor and ligand sorting. (Bottom) In the vulval precursor cells, AP-1, ARF-1, and AGEF-1 inhibit apical membrane localization of LIN-12/Notch required for secondary vulval fate (2°) induction. This antagonism is possibly due to their basolateral sorting of LIN-12/Notch. Considering AP-1, ARF-1 and AGEF-1’s established inhibition of primary vulval fate (1°) induction, it is also possible that they promote induction of the tertiary non-vulval fate (3°).

AGEF-1/ARF-1/AP-1 promotes LIN-12/Notch signaling during the AC/VU decision in the somatic gonad. This was evidenced by the *agef-1(vh4), arf-1(RNAi)* and *unc-101(sy108)* restoring the AC fate in the *lin-12(n302)* weak hypermorph, as well as reduced LIN-12 biosensor activation and decreased expression of LIN-12/Notch and the LAG-2/DSL ligand in *unc-101(sy108)*. However, *unc-101(sy108)* did not restore the AC fate in the *lin-12(n950)* strong hypermorph, suggesting that the signal may be too strong to be suppressed by a partial reduction in LIN-12 and LAG-2 membrane localization. Thus, AP-1 is a non-essential positive regulator of LIN-12/Notch signaling during the AC/VU decision.

Considering its canonical trafficking function [3, 38], AP-1 likely sorts the LIN-12/Notch and the LAG-2/DSL ligand between the TGN, recycling endosomes and the plasma membrane, such that their reduced expressions upon loss of AP-1 may reflect an increase in their degradation. Knockdown of AP-1 γ1 subunit in HeLa cells reduced the plasma membrane expression of >900 proteins suggesting a wide-spread role in protein trafficking to the plasma membrane [39]. Both LIN-12/Notch and LAG-2/DSL have a putative tyrosine and candidate dileucine based AP-1 binding sites suggesting that they could direct targets of AP-1 trafficking (Fig. S4) [40, 41]. Mutating these sites in future studies might provide more information on how they are trafficked but could be confounded by the fact that the same sites may be used by AP-2 and AP-3 adaptor complexes. While LIN-12/Notch and LAG-2/DSL levels were decreased we cannot rule out that AP-1 could be regulating the localization of other Notch signaling components such as the gamma secretase complex which also resides in the plasma membrane [42].

**Fig S4.**
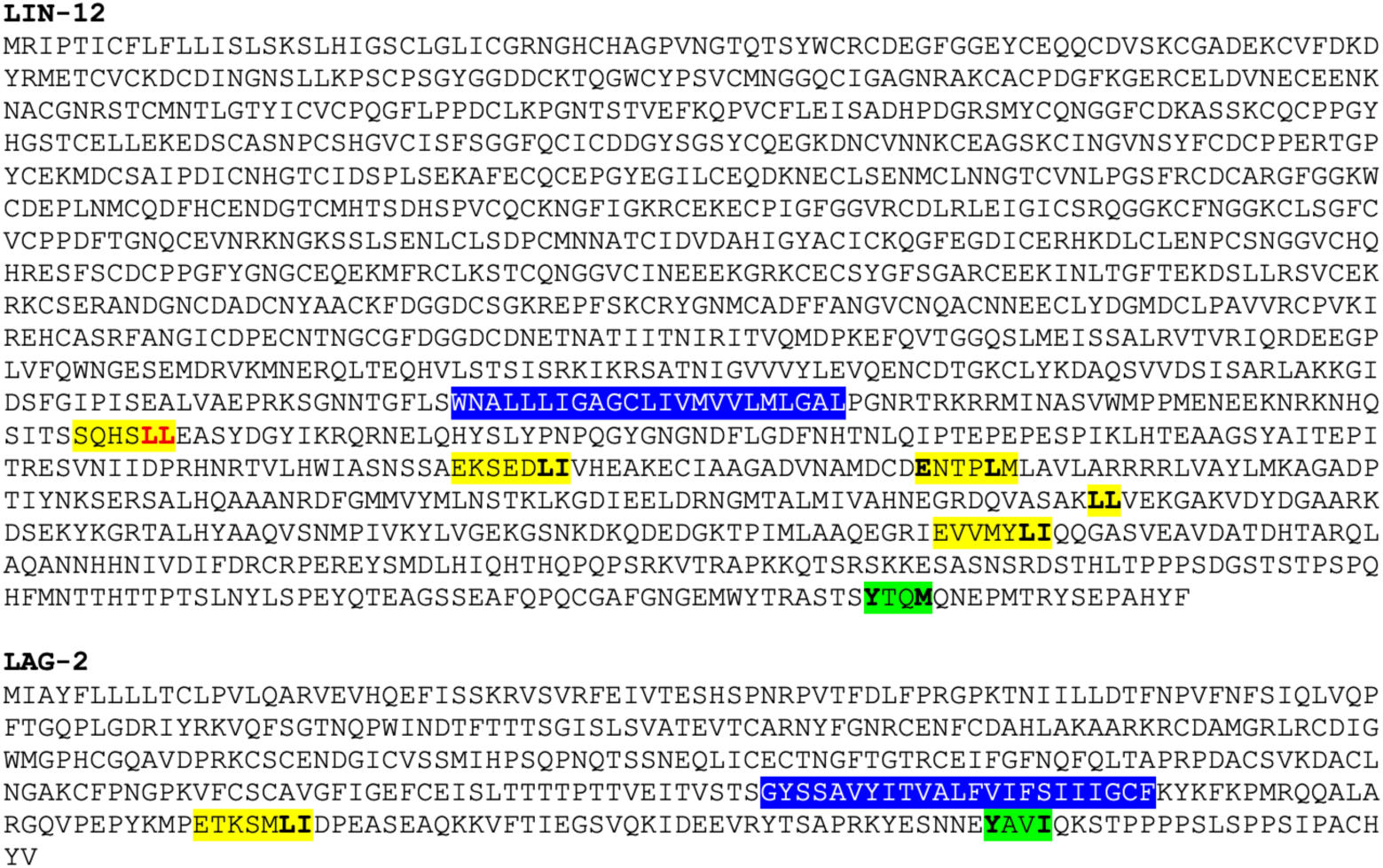
Candidate AP-1 sorting motifs in LIN-12 and LAG-2 cytoplasmic domains. LIN-12 and LAG-2 protein sequences. Transmembrane domains are highlighted in blue. Putative tyrosine-based sorting motifs (YXXM/I/L/V/F) are highlight in green. Potential dileucine sorting motifs (D/EXXXLL/I) are shown in yellow. The red dileucine motif was demonstrated to regulate LIN-12 localization in the VPCs [43, 44].

Despite promoting LIN-12/Notch signaling in the somatic gonad, we found that AGEF-1/ARF-1/AP-1 antagonizes signaling during vulva induction. This was evidenced by *agef-1(vh4), arf-1(RNAi)* and *unc-101(sy101)* inducing ectopic induction of secondary cell fates in the *lin-12(n302)* background, as well as increased LIN-12 biosensor activity and increased LIN-12 localization to the apical membrane of the presumptive secondary cells in *unc-101(sy108).* LIN-12/Notch is predominantly expressed on the apical membrane of the VPCs [43, 45], with its basolateral localization being insufficient for receptor activation [46]. Therefore, it is possible that AP-1 regulates LIN-12/Notch basolateral sorting to limit the levels on the apical membrane and *unc-101* mutants lead to LIN-12/Notch apical rerouting and subsequent signaling upregulation in the VPCs. Interestingly, a dileucine motif in the cytoplasmic domain of LIN-12 (Fig. S4, shown in red) was shown to regulate LIN-12 trafficking. Mutating this motif to alanines blocked endocytosis and led to accumulation at the apical membrane [43, 44] and thus could be a direct site of regulation by AP-1 and/or AP-2.

Interestingly, AP-1 differentially regulates Notch signaling in Drosophila. During eye development AP-1 promotes Notch signaling by regulating the trafficking of Scabrous [26]. Scabrous interacts with the extracellular domain of Notch where it can stabilize Notch at the plasma membrane. Loss of AP-1 results in intracellular accumulation of Scabrous and lysosomal degradation of Notch. In contrast, AP-1 antagonizes Notch signaling during sensory organ precursor cell specification. Loss of AP-1 caused phenotypes consistent with increased Notch activity, increased apical localization of the Notch activator Sanpodo as well as stabilization of Notch and Sanpodo at adherens junctions [25][26]. With no obvious homologs of Sanpodo and Scabrous in *C. elegans*, the differential regulation of LIN-12/Notch signaling might be due to other tissue specific regulators or factors.

Another explanation for the differences in regulation could be due to the VPCs being epithelial cells while the α cells are not. The presence of distinct apical and basal membranes poses more complex signaling pathways to sort proteins to the correct membrane compartments [47, 48]. AP-1 has been shown to differentially regulate protein trafficking to apical and basolateral membranes [38]. Since AP-1 promotes apical localization of LET-23/EGFR and basolateral localization of LIN-12/Notch, it is possible that different cargoes are trafficked differently by AP-1 with distinct coregulators. Alternatively, LET-23/EGFR or LIN-12/Notch could indirectly be regulated by AP-1. AP-1 has been shown to maintain apicobasal polarity in the intestinal cells and thus a general disruption of apicobasal polarity in the VPCs could result in mislocalization of LET-23/EGFR and LIN-12/Notch. We did not observe any obvious mislocalization of apical (PAR-6/Par6) or basolateral (LET-413/Scribble) polarity markers in the VPC daughters of *unc-101(sy101)* mutants (Fig. S5). However, LET-413/Scribble was unexpectedly apical in wild-type animals. Some apical mislocalization of LGL-1/Lethal Giant Larvae was seen in the grand daughters of the VPCs in *unc-101(sy108)* when combined with *arf-1(RNAi).* It is possible we did not assess the correct polarity markers or sufficiently reduce the activity of the AGEF-1/ARF-1/AP-1 pathway to cause visible differences. If AP-1 antagonizes LET-23/EGFR and LIN-12/Notch signaling indirectly by regulating cell polarity, then we expect that disrupting cell polarity in the VPCs might regulate signaling.

**Fig. S5.**
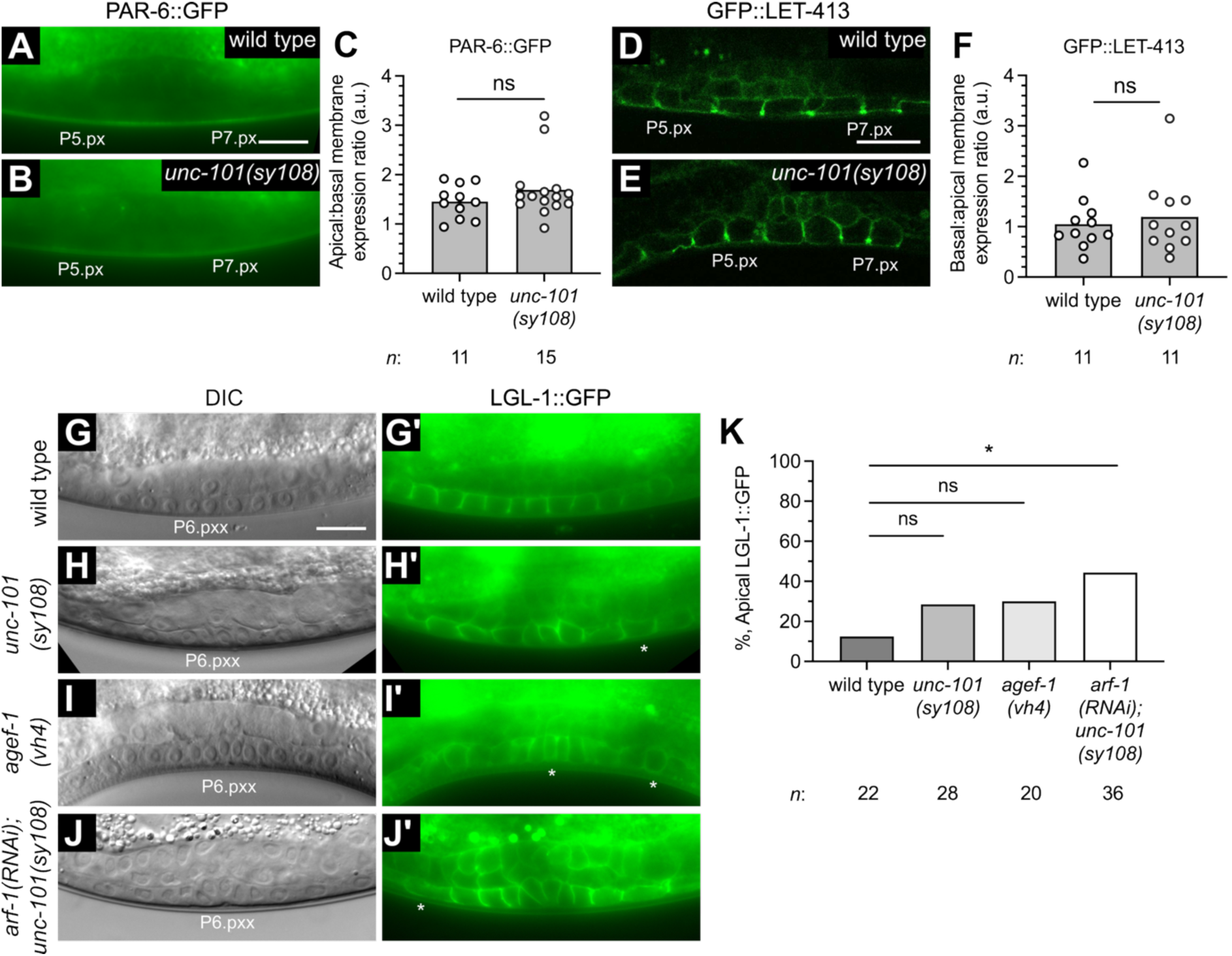
AP-1 does not significantly regulate localizations of polarity proteins PAR-6, LET-413 or LGL-1 in the VPC descendents. (A, B) Epifluorescent images of the apical polarity protein PAR-6::GFP, *xnIs3,* in the VPC daughters (Pn.px cells) of wild type and *unc-101(sy108)*. (C) Apical:basal VPC membrane expression ratios in wild type and *unc-101(sy108)*. Unpaired t-test. ns, not significant. (D, E) Confocal images of the basolateral/junctional polarity protein GFP::LET-413*, let-413(mib81),* in the VPC daughters of wild type and *unc-101(sy108)*. (F) Apical:basal VPC membrane expression ratios in wild type and *unc-101(sy108)*. Unpaired t-test. (G-J) Differential interference contrast (DIC) and epifluorescence images of basolateral protein LGL-1::GFP, *lgl-1(crk66)*, in the VPC granddaughters (Pn.pxx) of wild type, *unc-101(sy108)*, *agef-1(vh4)* and *arf-1(RNAi); unc-101(sy108)*. Asterisks denote VPC daughter cells with apically mislocalized LGL-1::GFP. (K) Percent animals with apical mislocalization of LGL-1::GFP in wild type, *unc-101(sy108)*, *agef-1(vh4)* and *arf-1(RNAi); unc-101(sy108)*. One-way ANOVA. ns, not significant, *P<0.05. Scale bars, 10µm.

The AGEF-1/ARF-1/AP-1 pathway is required for the fidelity of VPC patterning and may function to promote the tertiary non-vulval fate. Since this pathway is essential for viability, we used non-null allele of *agef-1* as well as null alleles of *arf-1* and *unc-101* which are partly redundant during vulva induction with their respective paralogs *arf-5* and *apm-1* [7, 16, 29]. Single mutants do not cause overt VPC induction phenotypes, but combined knockdown results in a Multivulva phenotype indicating that this pathway is critical for maintaining normal patterning of the VPCs [7, 9, 16]. Since the AGEF-1/ARF-1/AP-1 pathway was previously found to antagonize LET-23/EGFR signaling it was assumed that ectopic vulva induction was wholly contributed by increased LET-23/EGFR signaling. Our finding that AGEF-1/ARF-1/AP-1 pathway antagonizes LIN-12/Notch-mediated secondary cell fate induction implies that this pathway has a more widespread role in vulva induction inhibiting both the primary and secondary vulval fates. The tertiary non-vulval fate was previously thought to be passive uninduced fate. A recent study found that the RAP-2 GTPase and the MIG-15 kinase antagonizes both primary and secondary fates to promote the tertiary non-vulval fate [49]. Similarly, we propose that the AGEF-1/ARF/AP-1 trafficking could promote the non-vulval tertiary fate by limiting LET-23/EGFR and LIN-12/Notch signaling.

## Materials and methods

*Resources:* Wormbase (wormbase.org) and the Alliance of Genome Resources (alliancegenome.org) were essential in the planning and design of the research described [50, 51].

*Strain maintenance*: *C. elegans* strains were maintained on nematode growth medium agar plates with HB101 *E. coli* at 20℃ [52]. The Bristol strain N2 was used as wild type control. The strains used in this study are listed in Supplementary Table T1.

**Supplementary Table T1.**
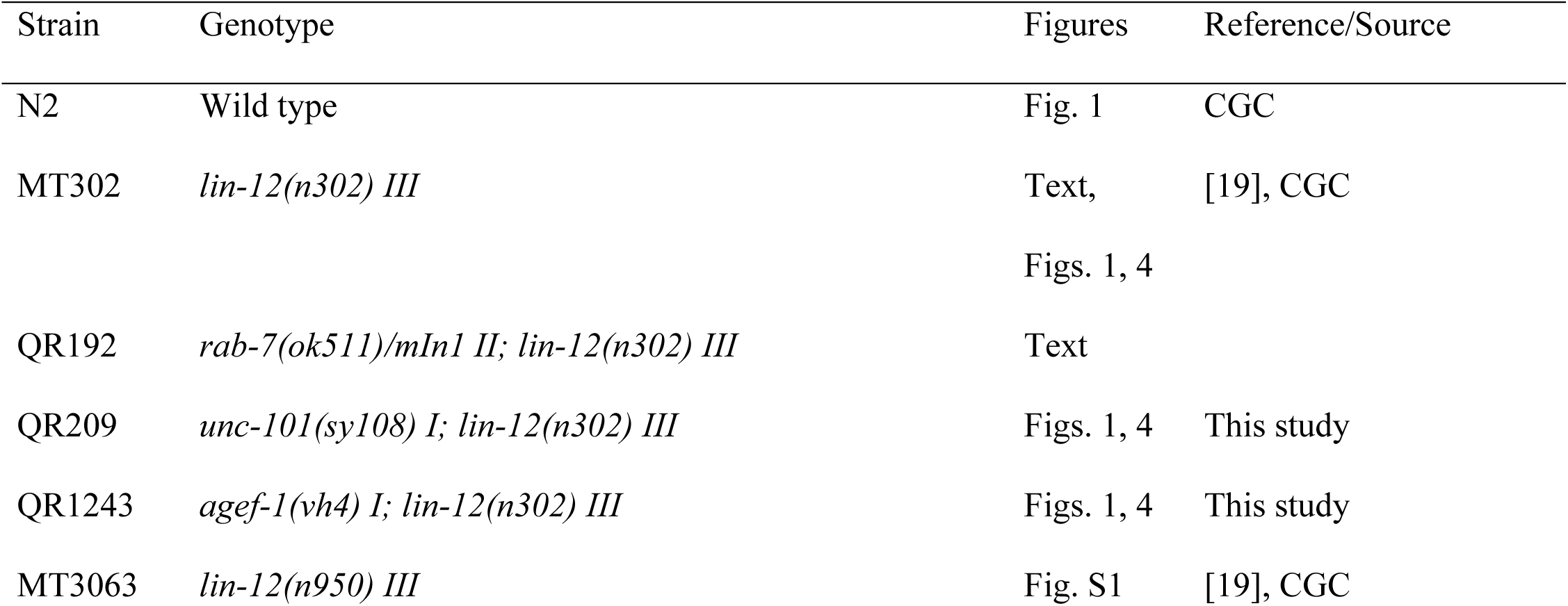
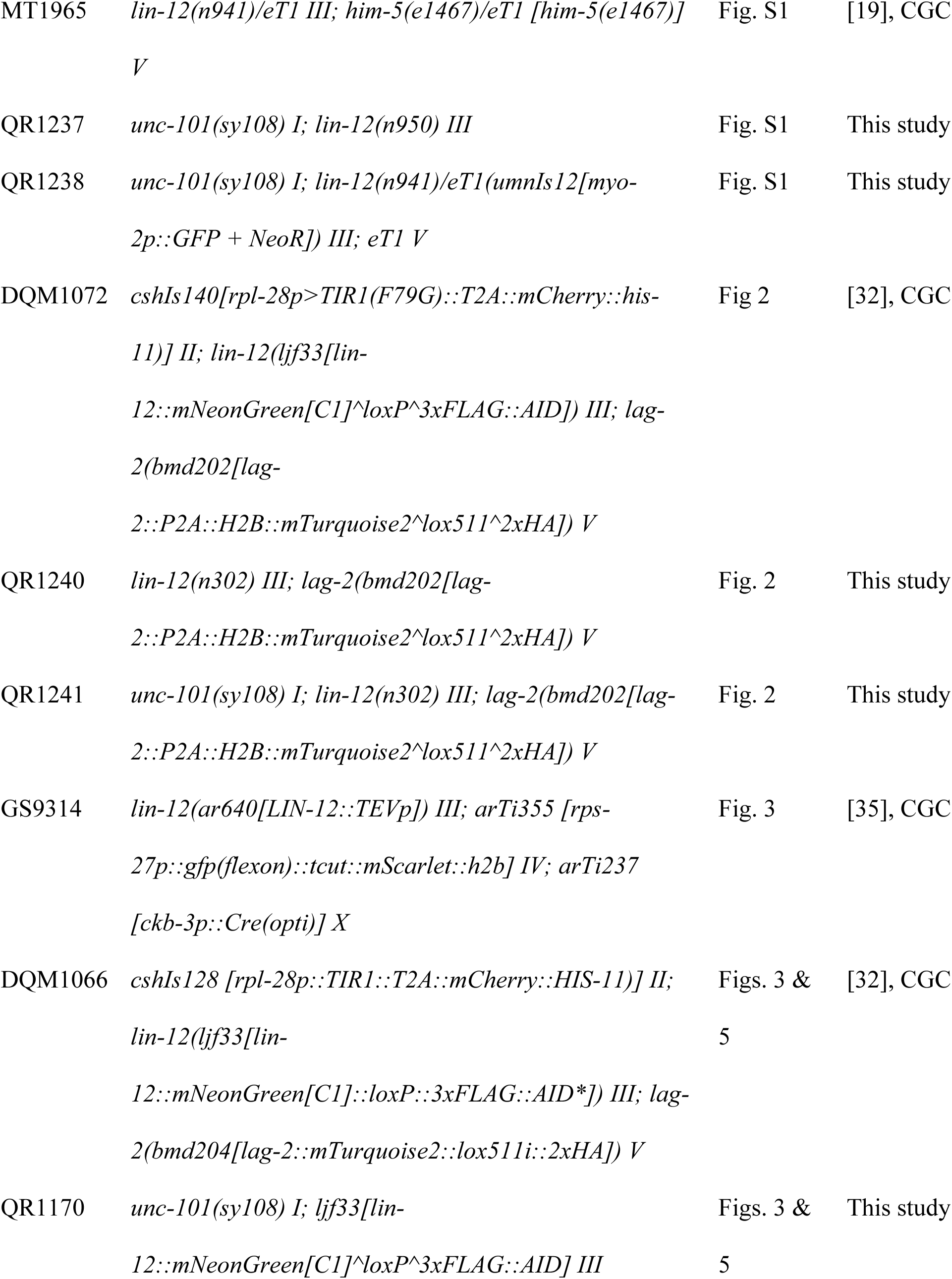
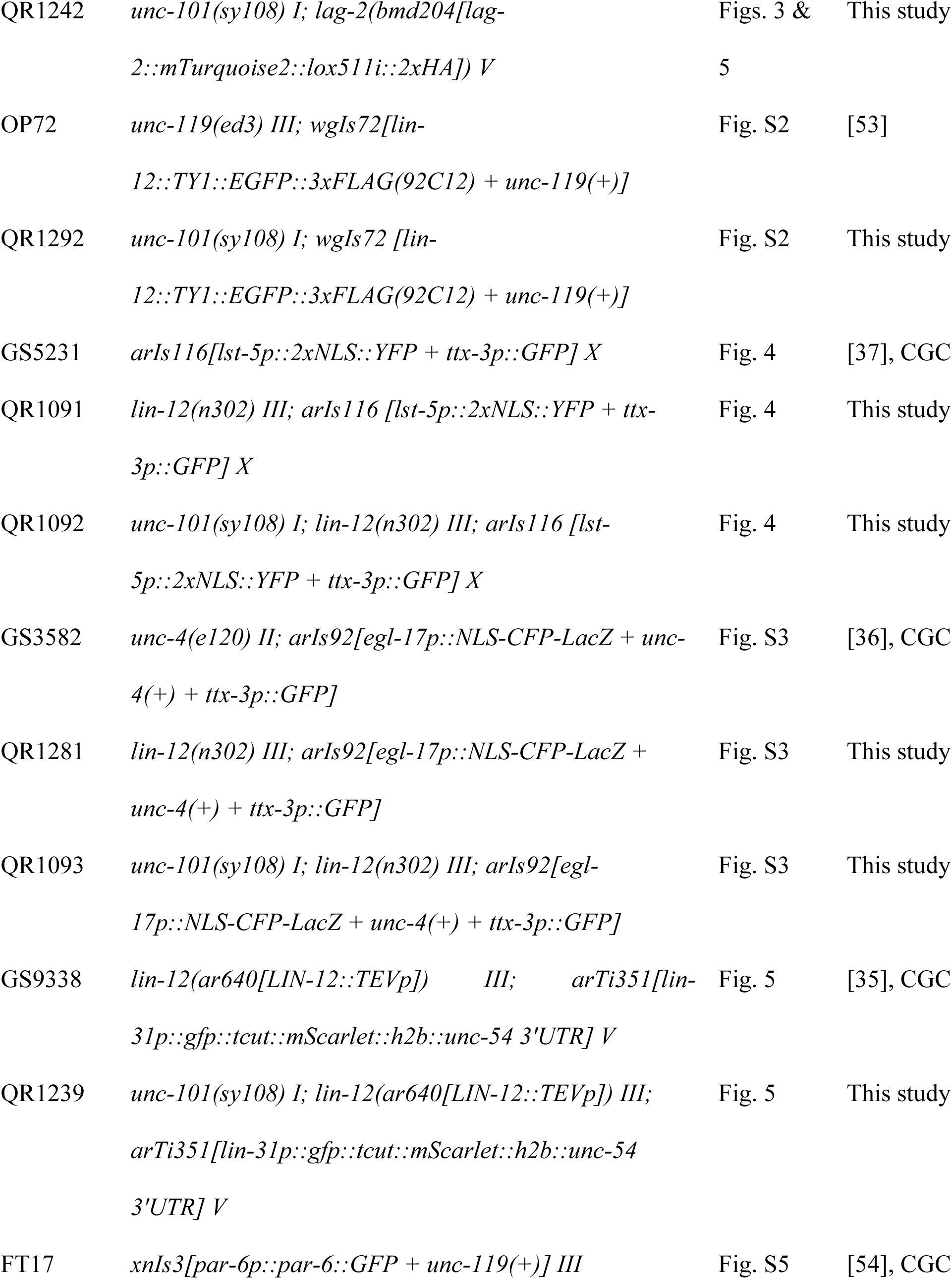
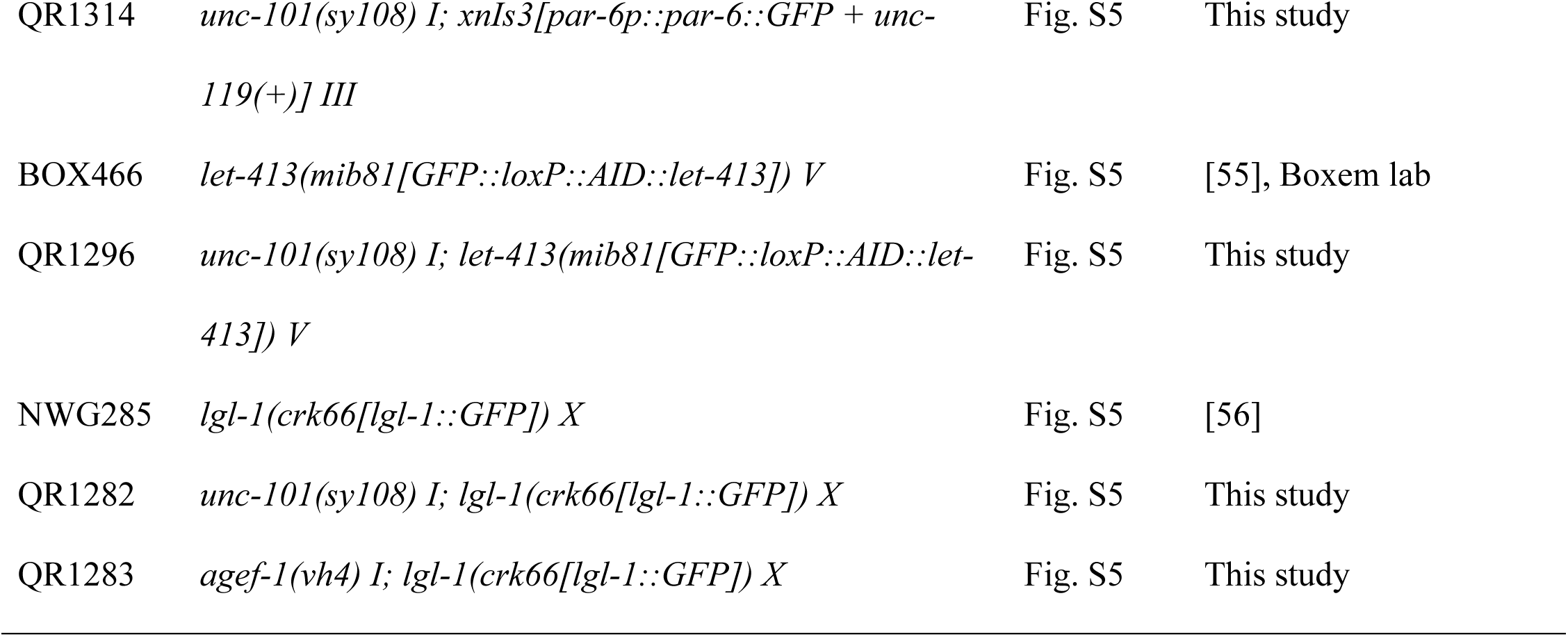
Strains used.

*RNA-mediated interference*: RNA-mediated interference (RNAi) by feeding was performed using the empty vector (L4440), *unc-101* (I-6G20), *arf-1* (III-3A13), and *agef-1* (I-6L24) clones of the Ahringer RNAi Library (Source BioScience, UK) [57–59]. To circumvent embryonic lethality caused by *arf-1(RNAi)* and *agef-1(RNAi)*, L1 larvae were synchronized as described [52], transferred onto seeded RNAi plates and imaged at various developmental stages. RNAi clones were verified by sequencing.

*Microscopy and image analysis*: Larvae were transferred to a 2% agarose pad in 10mM levamisole on a slide with a coverslip of either 0.13 to 0.16mm thickness for differential interference contrast (DIC) and epifluorescence images, or 0.16 to 0.19mm thickness for laser scanning confocal images. DIC and epifluorescence images were acquired on an Axio Imager A1 using the Axiocam 305 mono or AxioCam mRm camera with AxioVision software (Zeiss, Germany). Confocal images were acquired using an LSM780 laser-scanning confocal microscope with ZEN2010 software (Zeiss, Germany). To improve visibility, representative confocal images of the somatic gonad cells (Fig. 3) and the VPCs (Fig. 5) had their brightness and contrast modified equally across conditions. Confocal images of somatic gonad cells and the VPCs were analyzed using Fiji [60]. To quantify LIN-12/Notch signaling activity in the SALSA biosensor strains, nuclear selections of the maximum intensity projections RFP channel were used to measure mean intensity ratios of the RFP and GFP channels. To quantify LIN-12/Notch and LAG-2/DSL membrane expressions levels, membrane selections of the DIC channel were used to measure mean fluorescence intensity. To quantify LIN-12/Notch and LAG-2/DSL membrane expressions levels, membrane selections of the DIC channel were used to measure mean fluorescence intensity. To quantify expression ratios of apical and basolateral VPC membranes, the two intensity peaks corresponding to the apical and basolateral membranes were measured from a linear selection of the DIC channel through the P6.p membranes. Apical:basolateral ratios were measured using epifluorescent images of PAR-6::GFP animals, while in the case of GFP::LET-413 animals the single confocal Z-slice most focused on the nucleolus was analyzed. Frequencies of apical LGL-1::GFP expression were quantified in epifluorescent images of P5.pxx, P6.pxx and P7.pxx.

*VPC induction scoring*: Quantification of VPC induction was performed as described [61]. Three VPCs (P5.p-P7.p) are induced in wild-type hermaphrodites resulting in a VPC induction score of three. While induction occurs during the Pn.p stage, the VPC daughters (Pn.px) remain competent to be induced and thus when signaling is compromised it is possible for one of the two daughters to become induced resulting in a half induction. Vul animals have an induction score of 0 to 2.5 and Multivulva animals have an induction score of 3.5 to 6.

## Abbreviations

AC: Anchor Cell
AP: adaptor protein complex
DSL: delta serrate lag-2
EGFR: epidermal growth factor receptor
VPC: vulval precursor cell
VU: ventral uterine cell

## Acknowledgements

We thank Jung Hwa Seo for her technical assistance and support. We thank the Mike Boxem (Utrecht University), Iva Greenwald (Columbia University), Dave Matus (University of California, Berkley), and Jeremy Van Raamsdonk (McGill University) for their kind gifts of strains and reagents. Many strains were received from, or derived from, strains available at the Caenorhabditis Genetics Center (CGC), which is funded by NIH Office of Research Infrastructure Programs (P40 OD010440). Confocal imaging was performed at the Molecular Imaging Platform of the RI-MUHC. The RI-MUHC is supported in part by the Fonds de recherche du Québec−santé (FRQS).

## References

1. Farhan H, Rabouille C. Signalling to and from the secretory pathway. J Cell Sci. 2011;124(Pt 2):171–80. doi: 10.1242/jcs.076455. PubMed PMID: 21187344.

2. Sorkin A, von Zastrow M. Endocytosis and signalling: intertwining molecular networks. Nat Rev Mol Cell Biol. 2009;10(9):609–22. doi: 10.1038/nrm2748. PubMed PMID: 19696798; PubMed Central PMCID: PMCPMC2895425.

3. Bonifacino JS. Adaptor proteins involved in polarized sorting. J Cell Biol. 2014;204(1):7–17. Epub 2014/01/08. doi: 10.1083/jcb.201310021. PubMed PMID: 24395635; PubMed Central PMCID: PMCPMC3882786.

4. Nakatsu F, Hase K, Ohno H. The Role of the Clathrin Adaptor AP-1: Polarized Sorting and Beyond. Membranes (Basel). 2014;4(4):747–63. Epub 20141107. doi: 10.3390/membranes4040747. PubMed PMID: 25387275; PubMed Central PMCID: PMCPMC4289864.

5. Montpetit A, Cote S, Brustein E, Drouin CA, Lapointe L, Boudreau M, et al. Disruption of AP1S1, causing a novel neurocutaneous syndrome, perturbs development of the skin and spinal cord. PLoS Genet. 2008;4(12):e1000296. Epub 2008/12/06. doi: 10.1371/journal.pgen.1000296. PubMed PMID: 19057675; PubMed Central PMCID: PMCPMC2585812.

6. Alsaif HS, Al-Owain M, Barrios-Llerena ME, Gosadi G, Binamer Y, Devadason D, et al. Homozygous Loss-of-Function Mutations in AP1B1, Encoding Beta-1 Subunit of Adaptor-Related Protein Complex 1, Cause MEDNIK-like Syndrome. Am J Hum Genet. 2019;105(5):1016–22. Epub 2019/10/22. doi: 10.1016/j.ajhg.2019.09.020. PubMed PMID: 31630791; PubMed Central PMCID: PMCPMC6848991.

7. Shim J, Sternberg PW, Lee J. Distinct and redundant functions of mu1 medium chains of the AP-1 clathrin-associated protein complex in the nematode Caenorhabditis elegans. Mol Biol Cell. 2000;11(8):2743–56. Epub 2000/08/10. doi: 10.1091/mbc.11.8.2743. PubMed PMID: 10930467; PubMed Central PMCID: PMCPMC14953.

8. Boehm M, Bonifacino JS. Adaptins: the final recount. Mol Biol Cell. 2001;12(10):2907–20. doi: 10.1091/mbc.12.10.2907. PubMed PMID: 11598180; PubMed Central PMCID: PMCPMC60144.

9. Lee J, Jongeward GD, Sternberg PW. unc-101, a gene required for many aspects of Caenorhabditis elegans development and behavior, encodes a clathrin-associated protein. Genes Dev. 1994;8(1):60–73. doi: 10.1101/gad.8.1.60. PubMed PMID: 8288128.

10. Margeta MA, Wang GJ, Shen K. Clathrin adaptor AP-1 complex excludes multiple postsynaptic receptors from axons in C. elegans. Proc Natl Acad Sci U S A. 2009;106(5):1632–7. Epub 20090121. doi: 10.1073/pnas.0812078106. PubMed PMID: 19164532; PubMed Central PMCID: PMCPMC2635768.

11. Bae YK, Qin H, Knobel KM, Hu J, Rosenbaum JL, Barr MM. General and cell-type specific mechanisms target TRPP2/PKD-2 to cilia. Development. 2006;133(19):3859–70. Epub 20060830. doi: 10.1242/dev.02555. PubMed PMID: 16943275.

12. Dwyer ND, Adler CE, Crump JG, L’Etoile ND, Bargmann CI. Polarized dendritic transport and the AP-1 mu1 clathrin adaptor UNC-101 localize odorant receptors to olfactory cilia. Neuron. 2001;31(2):277–87. doi: 10.1016/s0896-6273(01)00361-0. PubMed PMID: 11502258.

13. Gillard G, Shafaq-Zadah M, Nicolle O, Damaj R, Pecreaux J, Michaux G. Control of E-cadherin apical localisation and morphogenesis by a SOAP-1/AP-1/clathrin pathway in C. elegans epidermal cells. Development. 2015;142(9):1684–94. Epub 20150409. doi: 10.1242/dev.118216. PubMed PMID: 25858456.

14. Feng S, Knodler A, Ren J, Zhang J, Zhang X, Hong Y, et al. A Rab8 guanine nucleotide exchange factor-effector interaction network regulates primary ciliogenesis. J Biol Chem. 2012;287(19):15602–9. Epub 20120319. doi: 10.1074/jbc.M111.333245. PubMed PMID: 22433857; PubMed Central PMCID: PMCPMC3346093.

15. Shafaq-Zadah M, Brocard L, Solari F, Michaux G. AP-1 is required for the maintenance of apico-basal polarity in the C. elegans intestine. Development. 2012;139(11):2061–70. Epub 20120425. doi: 10.1242/dev.076711. PubMed PMID: 22535414.

16. Skorobogata O, Escobar-Restrepo JM, Rocheleau CE. An AGEF-1/Arf GTPase/AP-1 ensemble antagonizes LET-23 EGFR basolateral localization and signaling during C. elegans vulva induction. PLoS Genet. 2014;10(10):e1004728. Epub 2014/10/21. doi: 10.1371/journal.pgen.1004728. PubMed PMID: 25329472; PubMed Central PMCID: PMCPMC4199573.

17. Schmid T, Hajnal A. Signal transduction during C. elegans vulval development: a NeverEnding story. Curr Opin Genet Dev. 2015;32:1–9. Epub 20150209. doi: 10.1016/j.gde.2015.01.006. PubMed PMID: 25677930.

18. Greenwald I. LIN-12/Notch signaling in C. elegans. WormBook. 2005:1–16. Epub 20050808. doi: 10.1895/wormbook.1.10.1. PubMed PMID: 18050403; PubMed Central PMCID: PMCPMC4781465.

19. Greenwald IS, Sternberg PW, Horvitz HR. The lin-12 locus specifies cell fates in Caenorhabditis elegans. Cell. 1983;34(2):435–44. doi: 10.1016/0092-8674(83)90377-x. PubMed PMID: 6616618.

20. Seydoux G, Greenwald I. Cell autonomy of lin-12 function in a cell fate decision in C. elegans. Cell. 1989;57(7):1237–45. doi: 10.1016/0092-8674(89)90060-3. PubMed PMID: 2736627.

21. Wilkinson HA, Fitzgerald K, Greenwald I. Reciprocal changes in expression of the receptor lin-12 and its ligand lag-2 prior to commitment in a C. elegans cell fate decision. Cell. 1994;79(7):1187–98. doi: 10.1016/0092-8674(94)90010-8. PubMed PMID: 8001154.

22. Hill RJ, Sternberg PW. The gene lin-3 encodes an inductive signal for vulval development in C. elegans. Nature. 1992;358(6386):470–6. doi: 10.1038/358470a0. PubMed PMID: 1641037.

23. Zand TP, Reiner DJ, Der CJ. Ras effector switching promotes divergent cell fates in C. elegans vulval patterning. Dev Cell. 2011;20(1):84–96. doi: 10.1016/j.devcel.2010.12.004. PubMed PMID: 21238927; PubMed Central PMCID: PMCPMC3028984.

24. Ferguson EL, Horvitz HR. Identification and characterization of 22 genes that affect the vulval cell lineages of the nematode Caenorhabditis elegans. Genetics. 1985;110(1):17–72. doi: 10.1093/genetics/110.1.17. PubMed PMID: 3996896; PubMed Central PMCID: PMCPMC1202554.

25. Benhra N, Lallet S, Cotton M, Le Bras S, Dussert A, Le Borgne R. AP-1 controls the trafficking of Notch and Sanpodo toward E-cadherin junctions in sensory organ precursors. Curr Biol. 2011;21(1):87–95. Epub 20101230. doi: 10.1016/j.cub.2010.12.010. PubMed PMID: 21194948.

26. Kametaka S, Kametaka A, Yonekura S, Haruta M, Takenoshita S, Goto S, et al. AP-1 clathrin adaptor and CG8538/Aftiphilin are involved in Notch signaling during eye development in Drosophila melanogaster. J Cell Sci. 2012;125(Pt 3):634–48. doi: 10.1242/jcs.090167. PubMed PMID: 22389401.

27. Skorobogata O, Rocheleau CE. RAB-7 antagonizes LET-23 EGFR signaling during vulva development in Caenorhabditis elegans. PLoS One. 2012;7(4):e36489. Epub 2012/05/05. doi: 10.1371/journal.pone.0036489. PubMed PMID: 22558469; PubMed Central PMCID: PMCPMC3340361.

28. Greenwald I, Seydoux G. Analysis of gain-of-function mutations of the lin-12 gene of Caenorhabditis elegans. Nature. 1990;346(6280):197–9. doi: 10.1038/346197a0. PubMed PMID: 2164160.

29. FitzPatrick C, Skorobogata O, Fazlollahi AM, Gauthier KD, Rocheleau CE. An activating mutation in AGEF-1, a putative Arf GEF, causes yolk extrusion from C. elegans embryos. bioRxiv. 2026. Epub 20260211. doi: 10.64898/2026.02.10.705107. PubMed PMID: 41727030; PubMed Central PMCID: PMCPMC12918894.

30. Newman AP, Sternberg PW. Coordinated morphogenesis of epithelia during development of the Caenorhabditis elegans uterine-vulval connection. Proc Natl Acad Sci U S A. 1996;93(18):9329–33. doi: 10.1073/pnas.93.18.9329. PubMed PMID: 8790329; PubMed Central PMCID: PMCPMC38427.

31. Newman AP, White JG, Sternberg PW. Morphogenesis of the C. elegans hermaphrodite uterus. Development. 1996;122(11):3617–26. doi: 10.1242/dev.122.11.3617. PubMed PMID: 8951077.

32. Medwig-Kinney TN, Sirota SS, Gibney TV, Pani AM, Matus DQ. An in vivo toolkit to visualize endogenous LAG-2/Delta and LIN-12/Notch signaling in C. elegans. MicroPubl Biol. 2022;2022. Epub 20220728. doi: 10.17912/micropub.biology.000602. PubMed PMID: 35966395; PubMed Central PMCID: PMCPMC9372767.

33. Kimble J. Alterations in cell lineage following laser ablation of cells in the somatic gonad of Caenorhabditis elegans. Dev Biol. 1981;87(2):286–300. doi: 10.1016/0012-1606(81)90152-4. PubMed PMID: 7286433.

34. Kimble J, Hirsh D. The postembryonic cell lineages of the hermaphrodite and male gonads in Caenorhabditis elegans. Dev Biol. 1979;70(2):396–417. doi: 10.1016/0012-1606(79)90035-6. PubMed PMID: 478167.

35. Shaffer JM, Greenwald I. SALSA, a genetically encoded biosensor for spatiotemporal quantification of Notch signal transduction in vivo. Dev Cell. 2022;57(7):930–44 e6. doi: 10.1016/j.devcel.2022.03.008. PubMed PMID: 35413239; PubMed Central PMCID: PMCPMC9473748.

36. Yoo AS, Bais C, Greenwald I. Crosstalk between the EGFR and LIN-12/Notch pathways in C. elegans vulval development. Science. 2004;303(5658):663–6. doi: 10.1126/science.1091639. PubMed PMID: 14752159.

37. Li J, Greenwald I. LIN-14 inhibition of LIN-12 contributes to precision and timing of C. elegans vulval fate patterning. Curr Biol. 2010;20(20):1875–9. Epub 20101014. doi: 10.1016/j.cub.2010.09.055. PubMed PMID: 20951046; PubMed Central PMCID: PMCPMC3322352.

38. Duncan MC. New directions for the clathrin adaptor AP-1 in cell biology and human disease. Curr Opin Cell Biol. 2022;76:102079. Epub 20220413. doi: 10.1016/j.ceb.2022.102079. PubMed PMID: 35429729; PubMed Central PMCID: PMCPMC9187608.

39. Wan C, Crisman L, Wang B, Tian Y, Wang S, Yang R, et al. AAGAB is an assembly chaperone regulating AP1 and AP2 clathrin adaptors. J Cell Sci. 2021;134(19). Epub 20211005. doi: 10.1242/jcs.258587. PubMed PMID: 34494650; PubMed Central PMCID: PMCPMC8520731.

40. Bonifacino JS, Traub LM. Signals for sorting of transmembrane proteins to endosomes and lysosomes. Annu Rev Biochem. 2003;72:395–447. Epub 20030306. doi: 10.1146/annurev.biochem.72.121801.161800. PubMed PMID: 12651740.

41. Robinson MS. Adaptable adaptors for coated vesicles. Trends Cell Biol. 2004;14(4):167–74. doi: 10.1016/j.tcb.2004.02.002. PubMed PMID: 15066634.

42. Goutte C. Genetics leads the way to the accomplices of presenilins. Dev Cell. 2002;3(1):6–7. doi: 10.1016/s1534-5807(02)00213-7. PubMed PMID: 12110162.

43. Shaye DD, Greenwald I. LIN-12/Notch trafficking and regulation of DSL ligand activity during vulval induction in Caenorhabditis elegans. Development. 2005;132(22):5081–92. Epub 20051019. doi: 10.1242/dev.02076. PubMed PMID: 16236769.

44. Shaye DD, Greenwald I. Endocytosis-mediated downregulation of LIN-12/Notch upon Ras activation in Caenorhabditis elegans. Nature. 2002;420(6916):686–90. doi: 10.1038/nature01234. PubMed PMID: 12478297.

45. Levitan D, Greenwald I. LIN-12 protein expression and localization during vulval development in C. elegans. Development. 1998;125(16):3101–9. doi: 10.1242/dev.125.16.3101. PubMed PMID: 9671583.

46. de Souza N, Vallier LG, Fares H, Greenwald I. SEL-2, the C. elegans neurobeachin/LRBA homolog, is a negative regulator of lin-12/Notch activity and affects endosomal traffic in polarized epithelial cells. Development. 2007;134(4):691–702. Epub 20070110. doi: 10.1242/dev.02767. PubMed PMID: 17215302.

47. Garcia-Castillo MD, Chinnapen DJ, Lencer WI. Membrane Transport across Polarized Epithelia. Cold Spring Harb Perspect Biol. 2017;9(9). Epub 20170901. doi: 10.1101/cshperspect.a027912. PubMed PMID: 28213463; PubMed Central PMCID: PMCPMC5585844.

48. Cao X, Surma MA, Simons K. Polarized sorting and trafficking in epithelial cells. Cell Res. 2012;22(5):793–805. Epub 20120424. doi: 10.1038/cr.2012.64. PubMed PMID: 22525333; PubMed Central PMCID: PMCPMC3343658.

49. Fakieh RA, Reiner DJ. RAP-2 and CNH-MAP4 Kinase MIG-15 confer resistance in bystander epithelium to cell-fate transformation by excess Ras or Notch activity. Proc Natl Acad Sci U S A. 2025;122(1):e2414321121. Epub 20241231. doi: 10.1073/pnas.2414321121. PubMed PMID: 39739816; PubMed Central PMCID: PMCPMC11725784.

50. Sternberg PW, Van Auken K, Wang Q, Wright A, Yook K, Zarowiecki M, et al. WormBase 2024: status and transitioning to Alliance infrastructure. Genetics. 2024;227(1). doi: 10.1093/genetics/iyae050. PubMed PMID: 38573366; PubMed Central PMCID: PMCPMC11075546.

51. Alliance of Genome Resources C. Updates to the Alliance of Genome Resources central infrastructure. Genetics. 2024;227(1). doi: 10.1093/genetics/iyae049. PubMed PMID: 38552170; PubMed Central PMCID: PMCPMC11075569.

52. Stiernagle T. Maintenance of C. elegans. WormBook. 2006:1–11. Epub 20060211. doi: 10.1895/wormbook.1.101.1. PubMed PMID: 18050451; PubMed Central PMCID: PMCPMC4781397.

53. Sarov M, Murray JI, Schanze K, Pozniakovski A, Niu W, Angermann K, et al. A genome-scale resource for in vivo tag-based protein function exploration in C. elegans. Cell. 2012;150(4):855–66. doi: 10.1016/j.cell.2012.08.001. PubMed PMID: 22901814; PubMed Central PMCID: PMCPMC3979301.

54. Anderson DC, Gill JS, Cinalli RM, Nance J. Polarization of the C. elegans embryo by RhoGAP-mediated exclusion of PAR-6 from cell contacts. Science. 2008;320(5884):1771–4. doi: 10.1126/science.1156063. PubMed PMID: 18583611; PubMed Central PMCID: PMCPMC2670547.

55. Riga A, Cravo J, Schmidt R, Pires HR, Castiglioni VG, van den Heuvel S, et al. Caenorhabditis elegans LET-413 Scribble is essential in the epidermis for growth, viability, and directional outgrowth of epithelial seam cells. PLoS Genet. 2021;17(10):e1009856. Epub 20211021. doi: 10.1371/journal.pgen.1009856. PubMed PMID: 34673778; PubMed Central PMCID: PMCPMC8570498.

56. Rodrigues NTL, Bland T, Borrego-Pinto J, Ng K, Hirani N, Gu Y, et al. SAIBR: a simple, platform-independent method for spectral autofluorescence correction. Development. 2022;149(14). Epub 20220714. doi: 10.1242/dev.200545. PubMed PMID: 35713287; PubMed Central PMCID: PMCPMC9445497.

57. Timmons L, Fire A. Specific interference by ingested dsRNA. Nature. 1998;395(6705):854. Epub 1998/11/06. doi: 10.1038/27579. PubMed PMID: 9804418.

58. Fraser AG, Kamath RS, Zipperlen P, Martinez-Campos M, Sohrmann M, Ahringer J. Functional genomic analysis of C. elegans chromosome I by systematic RNA interference. Nature. 2000;408(6810):325–30. Epub 2000/12/01. doi: 10.1038/35042517. PubMed PMID: 11099033.

59. Kamath RS, Fraser AG, Dong Y, Poulin G, Durbin R, Gotta M, et al. Systematic functional analysis of the Caenorhabditis elegans genome using RNAi. Nature. 2003;421(6920):231–7. Epub 2003/01/17. doi: 10.1038/nature01278. PubMed PMID: 12529635.

60. Schindelin J, Arganda-Carreras I, Frise E, Kaynig V, Longair M, Pietzsch T, et al. Fiji: an open-source platform for biological-image analysis. Nat Methods. 2012;9(7):676–82. Epub 20120628. doi: 10.1038/nmeth.2019. PubMed PMID: 22743772; PubMed Central PMCID: PMCPMC3855844.

61. Gauthier K, Rocheleau CE. C. elegans Vulva Induction: An In Vivo Model to Study Epidermal Growth Factor Receptor Signaling and Trafficking. Methods Mol Biol. 2017;1652:43–61 Epub 2017/08/10. doi: 10.1007/978-1-4939-7219-7_3. PubMed PMID: 28791633.

